# Attenuation of a DNA Cruciform by a Conserved Regulator Directs T3SS-1 mediated virulence in *Vibrio parahaemolyticus*

**DOI:** 10.1101/2022.03.07.483294

**Authors:** Landon J. Getz, Justin M. Brown, Lauren Sobot, Alexandra Chow, Jastina Mahendrarajah, Nikhil A. Thomas

## Abstract

Pathogenic *Vibrio* species account for 3-5 million annual life-threatening human infections. Virulence is driven by bacterial hemolysin and toxin gene expression often positively regulated by the winged helix-turn-helix (wHTH) HlyU transcriptional regulator family and silenced by Histone-like nucleoid structural protein (H-NS). In the case of *Vibrio parahaemolyticus*, HlyU is required for virulence gene expression associated with Type 3 Secretion System-1 (T3SS-1) although its mechanism of action is not understood. Here, we provide evidence for DNA cruciform attenuation mediated by HlyU binding to support concomitant virulence gene expression. Genetic and biochemical experiments revealed that upon HlyU mediated DNA cruciform attenuation, an intergenic cryptic promoter became accessible allowing for *exsA* mRNA expression and initiation of an ExsA autoactivation feedback loop at a separate ExsA-dependent promoter. Using a heterologous *E. coli* expression system, we reconstituted the dual promoter elements which revealed that HlyU binding and DNA cruciform attenuation were strictly required to initiate the ExsA autoactivation loop. The data indicate that HlyU acts to attenuate a transcriptional repressive DNA cruciform to support T3SS-1 virulence gene expression and reveals a non-canonical extricating gene regulation mechanism in pathogenic *Vibrio* species.

## Introduction

Pathogenic *Vibrio* species cause millions of life-threatening human infections annually^1^, as well as fatal infections in seafood organisms that contribute to major economic losses in aquaculture^2^. Primarily, *Vibrio* spp. are responsible for severe diarrheal disease (in the case of *Vibrio parahaemolyticus* and *Vibrio cholerae*), as well as wound infections causing necrotizing fasciitis (*Vibrio vulnificus).* Both *V. parahaemolyticus* and *V. cholerae* employ a complex array of pathogenicity factors, including pore forming toxins and secretion systems, to generate disease in a host^1^.

These pathogens possess DNA binding proteins that either repress or activate virulence gene expression, many of which alter DNA conformation^3–9^. Specifically, the HlyU family of winged helix-turn-helix (wHTH) DNA-binding proteins positively regulate a subset of *Vibrio* spp. virulence genes^9–12^. HlyU proteins dimerize and interact with consecutive major grooves of the double-stranded DNA helix and are modeled to induce variably angled DNA bends at intergenic regions^13, 14^. Small drug HlyU inhibitors have been discovered^15, 16^ and some have been shown to limit disease in animal models of infection^16^. The exact mechanism underlying HlyU positive gene regulation remains unknown. In the cases of *V. vulnificus* and *Vibrio anguillarum,* competitive DNA binding studies have implicated HlyU in overcoming virulence gene silencing mediated by Histone-like Nucleoid-Structuring protein (H-NS)^17, 18^. Additional genetic evidence in support of this regulatory paradigm was later found in *V. parahaemolyticus*, where *hlyU* was no longer required to support T3SS-1 activity in the absence of *hns*^9^. Moreover in *V. cholerae,* a parallel example between the wHTH ToxT transcriptional activator and H-NS repression has been studied in context of *ctxAB* (cholera toxin) expression^19, 20^. How transcriptional regulators like HlyU and ToxT impact H-NS repression is not completely understood and various models for de-repression of H-NS have been proposed^19, 21, 22^. In many cases, H-NS DNA binding sites do not directly overlap with those of those of the transcriptional regulators raising the possibility of DNA topology or protein-DNA interactions that function at a distance to impact gene expression. For numerous pathogens, H-NS binding to DNA acts to constrain localized DNA supercoiling often leading to dramatic changes in DNA topology such as looping, hairpins, and cruciforms^23–25^. In some cases, the altered DNA topology negatively impacts promoter accessibility for key transcriptional initiation events^26^. Pathogens must therefore possess a variety of mechanisms to overcome these repressive effects on virulence genes^27–31^.

In *Vibrio parahaemolyticus*, we previously discovered the *exsBA* intergenic region as an HlyU binding site, which contains inverted repeat sequences separated by A/T rich DNA^9^. Such DNA nucleotide arrangements are known to form DNA cruciform (or 4-way) structures under certain energetic provisions^24, 32, 33^. DNA cruciform structures are dynamic non-β-DNA structures found in all domains of life and are implicated in repressing gene expression ^24, 34^, replication initiation in bacteriophages and plasmids^32, 35^ and occasionally as DNA recombination intermediates^36^. The formation of DNA cruciform structures typically requires high superhelical density – as a mechanism of energy input into the DNA helix – constrained by negative supercoiling which is then relieved by contextual extrusion of DNA strands^37^. While DNA looping and bending mechanisms have been well characterized in bacterial genetic processes, functional DNA cruciform examples in bacterial chromosomes are rare and there are no examples linked to virulence gene regulation. Notably, protein crystallization studies have implicated HlyU binding to bent dsDNA^14, 38^, however no HlyU-DNA co-crystal has been solved to date and the functional genetic outcomes of this have not been investigated.

In this study, we set out to investigate the mechanism of HlyU function in context of *exsA* gene expression and subsequent T3SS-1 gene regulation. *V. parahaemolyticus* Δ*hlyU* mutants are known to be significantly impaired for *exsA* gene expression and exhibit reduced cytotoxicity during infection^9^. Conversely, *hns* null mutants exhibit significantly enhanced and de-regulated expression of *exsA*^9, 39, 40^. HlyU and H-NS binding sites within the *exsA* promoter region do not overlap and are separated by at least 100 base pairs^9, 40^ making competitive DNA binding models challenging to reconcile. Considering the nature and propensity of A/T rich supercoiled DNA to form cruciforms, we hypothesized that a DNA cruciform structure is involved in regulation of *exsA* expression. We present data that implicates HlyU in attenuating a transcriptionally repressive DNA cruciform leading to *V. parahaemolyticus* T3SS-1 mediated virulence.

## Results

### in silico identification of cruciform forming loci in Vibrio species

Extensive genomic analyses have revealed that potential cruciform forming DNA sequences are found to be significantly enriched at the 3’ end of genetic operons, often within intergenic regions and near transcriptional promoters^41, 42^. To investigate DNA sequences capable of forming cruciform structures within the *V. parahaemolyticus exsBA* intergenic region, we undertook an *in silico* approach using Palindrome Analyser^43^. We allowed for a maximum of one sequence mismatch in the base-paired cruciform stem DNA and excluded cruciforms with stem sequences which contained less than six nucleotides. Furthermore, DNA cruciforms with looped-DNA which are spaced by less than ten nucleotides were also excluded. These criteria were chosen to identify DNA sequences which together supported the annealing constraints for both DNA cruciform formation and HlyU binding^9–12^.

21 cruciform sequences were identified within the ∼650 nucleotide *exsBA* intergenic region which met our criteria for cruciform potential (table S1). Critically, these putative cruciform structures all had positive predicted *ΔG* values, which is consistent with an energetic supercoiling input requirement for cruciform formation. Three cruciform structures are clustered around the HlyU binding site and have similar *ΔG* values between 12.94 to 14.14 (table S1).

We also investigated if HlyU binding sites from other *Vibrio* spp. have cruciform forming potential. For all the *Vibrio* spp. evaluated, the HlyU binding site was in proximity or overlapped with a putative DNA cruciform structure (tables S2-4). These data indicate that DNA sequence features required for cruciform formation are present at a variety of intergenic HlyU binding sites in different *Vibrio* spp. The observations align with the known enrichment of cruciform forming DNA sequences at specific intergenic DNA locations^41^. In all cases, the respective DNA sequences exhibit A/T rich DNA (>90%) which aligns with the commonly described conditions that favor cruciform formation *in viv*o.^33, 35, 37, 44^

### Identification and Sequence Mapping of Cruciform Structures

To functionally evaluate the *Vibrio parahaemolyticus exsBA* intergenic region for cruciforms, we built on a previously developed T7 endonuclease assay with supercoiled plasmid DNA substrates^45^. Here, T7 endonuclease cleaves DNA cruciform structures in a sequential two-step process to produce a double-strand DNA cleavage and linearize plasmid DNA. A pUC(A/T) plasmid with an engineered stable cruciform served as a positive control for T7 endonuclease cleavage^45^ and empty pBluescript acted as a negative control (fig. 1a). As expected, treatment of pUC(A/T) with T7 endonuclease linearized the plasmid, while treatment of pBluescript did not (fig. 1b, pBS and pUC(A/T), see lane 4 for each condition).

**Fig. 1.**
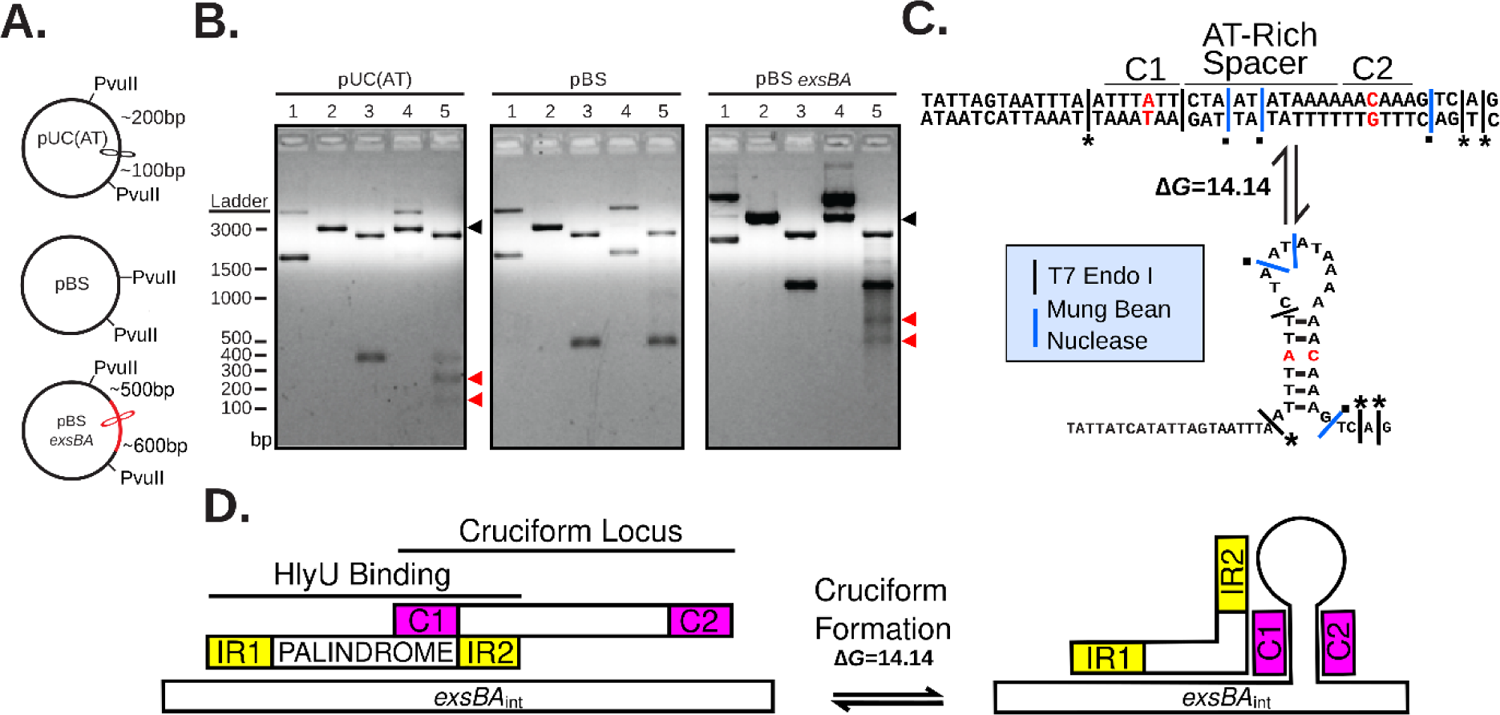
Identification of cruciform structures at the HlyU binding site in the *exsBA* locus of *V. parahaemolyticus.* (A) Schematic diagram of plasmid constructs used for T7 endonuclease I digestion analyses. Expected restriction fragment sizes and relative location of PvuII and cruciform sites are identified. The red line and cruciform structure indicate the cloned *exsBA* DNA region. (B) Restriction digestion of plasmid DNA visualized by agarose gel electrophoresis. 1-undigested plasmid, 2-linearized plasmid, 3-PvuII, 4-T7 Endonuclease, 5-T7 Endonuclease followed by PvuII. T7 Endonuclease targets cruciform structures to cause double-strand break in a two-step process. PvuII was used as it flanks the cloned DNA and allows for restriction mapping. The black arrowhead indicates linearized DNA for the respective cruciform forming plasmid constructs (compare lanes 2 and 4). Complete DNA cleavage by T7 endonuclease is rare due to the two step cut process (initial nicking) and dissipation of supercoiling in the intermediate stage, explaining the enriched slower migrating nicked DNA (lane 4, left and right panels). The red arrowheads in lane 5 of each panel indicate DNA cleavage fragments generated by initial cruciform cleavage by T7 endonuclease. pBluescript and pUC(AT) are negative and positive controls respectively. This experiment was repeated at least three times with representative data shown. (C) Schematic depicting cleavage results of the cruciform sequence mapping assay for the *exsBA* intergenic region in *V. parahaemolyticus.* Red nucleotides indicate stem mismatches in the cruciform sequence. The asterisks and black squares indicate cut sites that are consistent with the known specificities of T7 endonuclease and mung bean nuclease respectively. Cleavage data was obtained from multiple independent cloned DNA fragments (see methods). (D) Schematic diagram of the genomic locations of HlyU binding and the identified cruciform structure. Yellow rectangles depict inverted repeats (IR1 and IR2) that flank a palindromic DNA element that constitutes the HlyU binding site within the *exsBA* intergenic region. Cruciform stem DNA sequences (C1 and C2) are indicated by magenta rectangles. Only one ‘side’ of the stem-loop cruciform is shown in panels C and D for presentation purposes.

The *exsBA* intergenic region of *Vibrio parahaemolyticus* was cloned into a pBluescript vector (fig. 1a) and assayed by digestion with either T7 endonuclease alone or in combination with other restriction digests (fig. 1b). When recombinant pBluescript containing the *exsBA* intergenic region was treated with T7 endonuclease, dsDNA cleavage was consistently observed as evidenced by enrichment of DNA that migrated similarly to linearized DNA (fig. 1b, lane 4). This suggested the existence of a DNA cruciform within the *exsBA* intergenic region.

Initial T7 endonuclease digestion of the pUC(A/T) and *exsBA* constructs followed by sequential PvuII digestion provided additional DNA cleavage products that allowed approximation of cruciform position by DNA fragment sizes (fig. 1b, lane 5, fig. S1). As expected, uniform, cruciform-associated cleavage products of approximately 200 and 100 bp were consistently observed for pUC(A/T). For the *exsBA* construct DNA cleavage products appearing ∼600 and ∼450bp in size were observed. Importantly, these *exsBA* associated DNA cleavage products overlap the HlyU binding site^9^ and multiple predicted cruciform structures identified by Palindrome Analyser (table S1).

To further investigate the intergenic *exsBA* localized cruciform structure, we precisely sequence mapped the relevant T7 endonuclease cut sites (see methods). Here, we found compelling evidence for one of the cruciform structures identified by Palindrome Analyser (fig. 1c, table S1). Critically, three *exsBA* T7 endonuclease cut sites precisely mapped to the base of the cruciform stem, which is consistent with model synthetic cruciform cleavage studies (fig. 1c)^46^. Moreover, mung bean nuclease digestion assays efficiently localized single stranded DNA likely associated with ‘loop’ structures of the DNA cruciform (fig. 1c)^35^. These data indicate that a stable cruciform structure exists at a DNA locus that overlaps a HlyU binding site in *V. parahaemolyticus* (fig. 1d).

To further explore the cruciform forming capacity of HlyU binding sites in other *Vibrio* species, we subjected HlyU binding intergenic regions from *V. cholerae (tlh-hlyA – 1104bp), V. vulnificus (rtxA1* operon region *– 362bp)* and *V. anguillarum (plp-vah – 493bp)* to the T7 endonuclease and sequential cleavage experiments (fig. S1). T7 endonuclease cleavage produced linearization of the intergenic regions from *V. cholerae* and *V. vulnificus,* and unique digestion products were observed on sequential digest with PvuII (fig. S1d, e). However, no significant cleavage from either the sequential digest or the T7 endonuclease digestion alone was observed for the intergenic fragment from *V. anguillarum* (fig. S1f). A summary of cruciform location and cleavage data is presented in supplementary figure 2. These experiments provide evidence that cruciform structures may form at the site of HlyU binding in a variety of *Vibrio* spp.

**Fig. 2.**
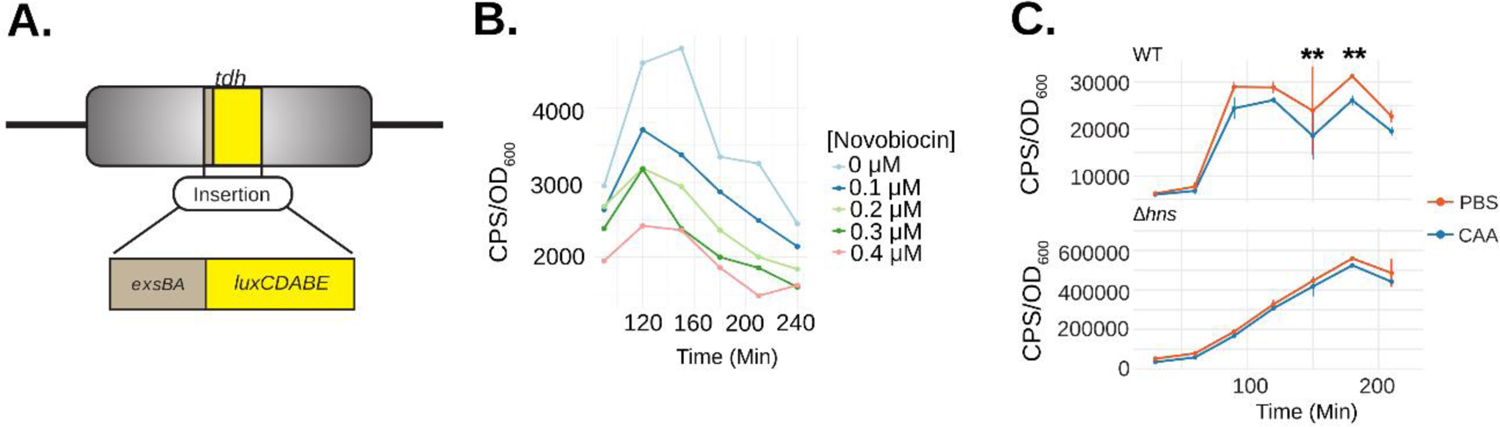
DNA supercoiling and cruciform formation within the *exsBA* genetic locus regulate *exsA* promoter activity. (A) Schematic depicting the *exsBA* intergenic-*luxCDABE* luciferase transcriptional reporter fusion integrated into the *V. parahaemolyticus tdh* chromosome locus to allow expression of the luciferase cassette (8). (B) *V. parahaemolyticus* harboring the *exsBA*-*luxCDABE* reporter (depicted in panel A) was treated with increasing concentrations of novobiocin and *exsA* promoter activity was quantitatively measured based on *in vivo* real-time light emission. (C) Wildtype *V. parahaemolyticus* and Δ*hns* strains harbouring the integrated *exsBA*-*luxCDABE* transcriptional reporter were treated with either phosphate buffered saline (PBS) or 40nM chloroacetylaldehyde (CAA). Light emission was measured as counts per second (CPS) every 30 minutes, along with OD600, for a total of 4 hours. Statistical significance was determined by multiple unpaired t-tests, ** p<0.01, n=2 and data were visualized using R and the ggplot2 package.

### Evidence for exsBA genetic locus DNA supercoiling and cruciform formation within V. parahaemolyticus cells

Next, we set out to find evidence for DNA cruciform formation on the *V. parahaemolyticus* chromosome within the *exsBA* intergenic region. We considered the possibility that the *exsBA* genetic locus is subject to DNA supercoiling, as DNA cruciform formation generally results from excessive torsional forces caused by DNA supercoiling. To investigate this possibility, we measured promoter activity for a chromosomally integrated *exsA* promoter fused to a *luxCDABE* reporter^9^ (fig. 2a). Bacteria were grown in the presence of increasing sublethal doses of novobiocin [0-0.4μM], which acts as a DNA gyrase inhibitor and potently reduces DNA supercoiling^47^. It was observed that *exsA* promoter activity was reduced in a dose dependent manner upon increasing concentration of novobiocin (fig. 2b). These data indicate that the *exsBA* intergenic region is subject to DNA supercoiling which impacts on transcriptional activity.

Next, we set out to investigate DNA cruciform formation within the *exsBA* intergenic region on chromosomal DNA. We employed a pulse-chase chemical treatment of cells with chloroacetaldehyde (CAA) which is known to interact with single-stranded DNA and unpaired bases to form DNA ethanoadducts^48^. We hypothesized that if a DNA cruciform exists at this locus in a proportion of cells, CAA treatment would reduce overall *exsA* promoter activity and thus negatively impact the expression of a transcriptional luciferase reporter.

The experiments were performed using an initial pulse of 40nM CAA which was determined to not inhibit subsequent cell growth (fig. S3). DNA ethenoadduct formation via CAA modification was performed in wildtype and *Δhns* strains containing an *exsA* promoter-*luxCDABE* reporter integrated into the chromosome of each strain to permit cis-regulation by DNA binding proteins, including HlyU and H-NS (fig. 2a). We reasoned that the *Δhns* mutant, owing to its lack of H-NS containment of DNA negative supercoiling, should be deficient for DNA cruciform structures, and generally less affected for *exsA* promoter activity. *in situ* real time *exsA* promoter activity measurements consistently revealed that the CAA-treated wildtype reporter strain exhibited significantly reduced *exsA* promoter activity when compared to untreated cells at 150 and 180 minutes (fig. 2c). In contrast, CAA treatment of the *Δhns* reporter strain did not exhibit a difference at these same time points using a 2-way ANOVA with Bonferroni multiple test correction (fig. 2c). These data suggest that CAA mediated ethanoadduct modification of unpaired DNA bases reduced transcriptional activity of the luciferase reporter. Moreover, H-NS containment of DNA supercoiling may play a role in the formation of the unpaired DNA within a cruciform structure. Importantly, these data are consistent with the well-established requirement for DNA supercoiling towards cruciform formation *in vivo*^33, 37^

### Inverted repeat and palindrome sequence elements contribute to HlyU binding

The HlyU binding site upstream of *exsA* in *V. parahaemolyticus* consists of a 55bp DNA stretch^9^ that partially overlaps the cruciform-forming locus (fig. 1d). We sought to investigate the DNA sequence within this 55bp region that is necessary for HlyU binding using electrophoretic mobility shift assays (EMSA). For these experiments, dsDNA fragments spanning 55bp were formed by annealing two single-stranded complementary oligos (see methods) and incubated with purified his-tagged HlyU (fig. S4). HlyU was able to bind to the wildtype DNA sequence consistently producing two shifted DNA species in EMSA experiments (denoted as S1 and S2, fig. 3a,c). Scrambling the palindrome sequence <u>ATTTAATTTA</u> between the perfect inverted repeats (ATATTAG and CTAATAT) to form mutant PAL1 abrogated HlyU binding to the DNA target (fig. 3a,b). This provided evidence that the palindrome sequence is necessary for HlyU interaction with DNA.

**Fig. 3.**
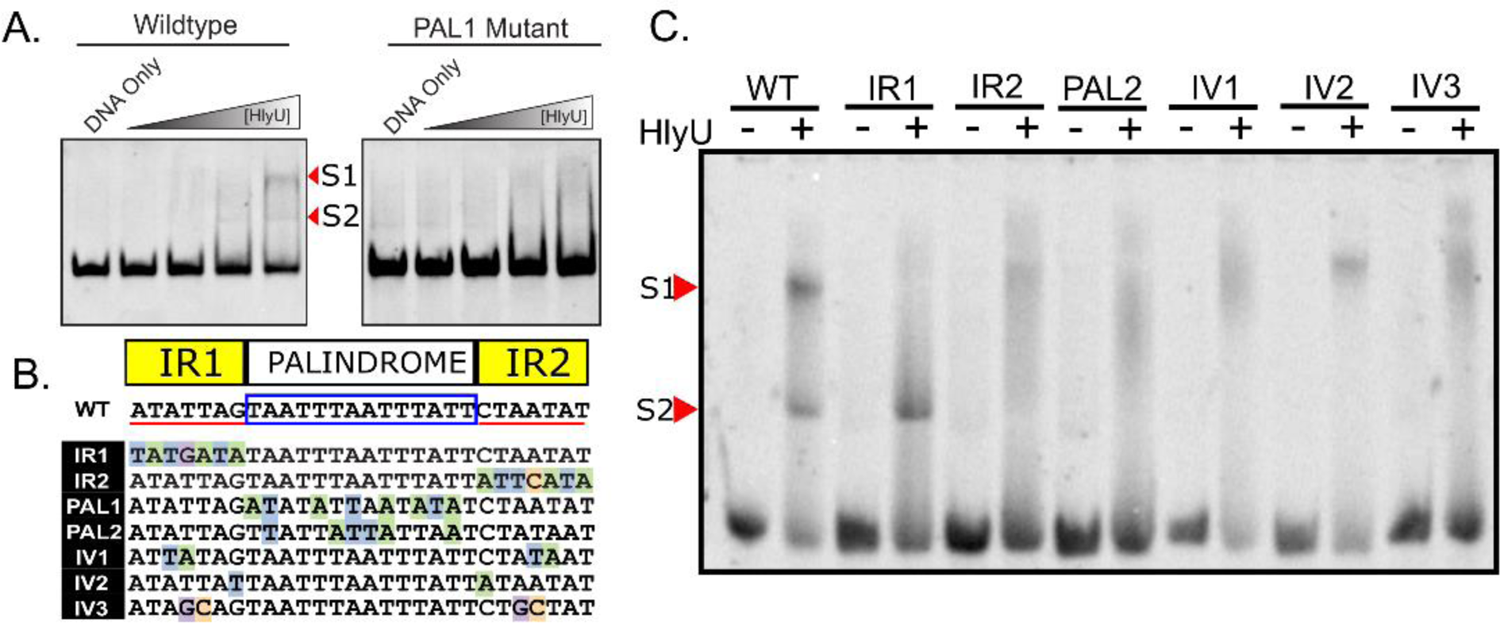
Mutational analysis identifies nucleotide elements important for HlyU interaction with DNA near the *exsBA* cruciform forming locus. (A) Oligonucleotides constituting a wildtype *exsBA* HlyU binding site, as well as a mutant in the palindromic region (PAL1), were annealed by sequential cooling and used as DNA targets for HlyU binding in electrophoretic mobilty shift assays (EMSA). Arrowheads refer to distinct bands that appear upon elevated amounts of purified HlyU. S1 and S2 indicate distinct shifted species in the EMSA assay. (B) Schematic of the specific mutant DNA sequences compared to wildtype (WT). Nucleotides with a coloured background identify differences from the wildtype sequence. Perfect inverted repeat DNA sequences are underlined in red separated by a boxed palindrome sequence. (C) Oligonucleotides of mutants described in (B) were annealed and subjected to EMSA with and without purified HlyU. Arrowheads refer to shifted HlyU-DNA complexes S1 and S2 All EMSA experiments were performed at least three times with representative data images shown.

To further explore this specific *exsBA* region, we generated a series of mutations in the inverted repeat sequences adjacent to the palindrome, as well as the palindromic sequence itself (fig. 3b). Every modification to these sequence elements impacted HlyU interaction with DNA by EMSA (fig. 3c) with one caveat. One of the inverted repeat mutations distal to *exsA* (IR1) partially bound HlyU producing only S2, the low molecular-weight shift species (fig. 3c). This is the only mutation that is adjacent (does not overlap) to the cruciform stem sequences (C1 and C2) previously identified (fig. 1d) (see discussion). Taken together, these data identify that both the inverted repeat sequences and the palindromic spacer are necessary DNA elements for efficient HlyU binding and that the cruciform-forming locus within the *exsBA* intergenic region partially overlaps with the site of HlyU binding.

### Mutations that alter HlyU binding negatively impact T3SS-1 activity in V. parahaemolyticus

The genetic and biochemical data suggested that HlyU binding to DNA attenuates a transcriptionally repressive cruciform leading to *exsA* expression. To address this further, we investigated the *V. parahaemolyticus* chromosomal intergenic *exsBA* locus by mutational analysis. Mutations were introduced onto the chromosome by allelic exchange and verified by DNA sequencing. The resulting mutant strains were compared to the parental strain, as well as T3SS-1 deficient Δ*hlyU* and Δ*vscN1* mutants^9, 49^ using protein secretion and host cell infection assays. T3SS-1 specific protein secretion was markedly reduced in each mutant, except for mutant IR1 which appeared slightly reduced, as detected by SDS-PAGE analysis (fig. 4a). The ability of these mutants to cause cytotoxicity towards HeLa cells during infection was significantly impaired compared to the parental strain (fig. 4b). The IR1 and IV2 mutants exhibited intermediate cytotoxicity and therefore retain intermediate T3SS-1 activity. HeLa cytotoxicity was restored in all the mutants by complementation with *exsA in trans* (table S5), which bypasses the effect of the chromosomal mutations. These *in vivo* analyses identify that mutations negatively impacting on HlyU binding produce a marked decrease in T3SS-1 activity as measured by secretion and infection (cytotoxicity) assays.

**Fig. 4.**
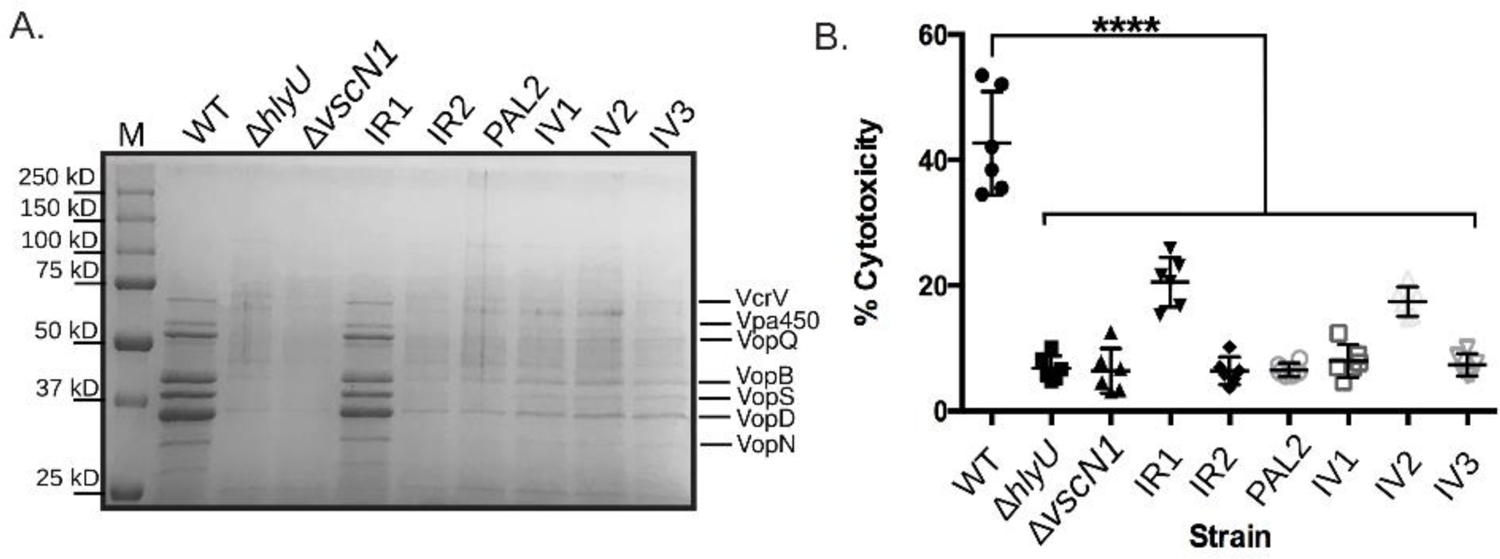
*V, parahaemolyticus* T3SS-1 protein secretion and host cell cytotoxicity are reduced in bacteria harbouring mutations at the HlyU binding site near the *exsBA* DNA cruciform forming locus. (A) Total secreted protein profiles derived from *V. parahaemolyticus* harbouring chromosomal mutations at the HlyU binding site and cruciform locus. The specific DNA mutations are listed in figure 3B and labeled accordingly for each gel lane. Proteins were visualized by Coomassie staining. M refers to protein standard. The indicated dominant protein species have been previously identified from WT using mass spectrometry^49^. The experiment was performed at least three times with a representative stained protein gel shown. (B) A LDH release assay from cultured HeLa cells was used to determine host cytotoxicity upon infection with the indicated *V. parahaemolyticus* strains. Statistical significance was determined by two-way ANOVA (against WT) using a Bonferroni test correction. **** p<0.001, n=3. Δ*hlyU* and Δ*vscN1* are known to be deficient for T3SS-1 activity and were included as comparative controls.

### HlyU binding attenuates a DNA cruciform in the exsBA intergenic region to support gene expression

Next, we set out to determine if HlyU binding to DNA serves to attenuate cruciform formation to support efficient gene expression. Informed by the cruciform mapping data (fig. 1) and EMSA data (fig. 3), we aimed to generate modified *exsBA* DNA constructs that could still form cruciforms but were unable or inefficient for HlyU binding. We reasoned that such DNA constructs could be generated given the intermediate phenotypes consistently observed for the IR1 and IV2 mutant bacterial strains (fig. 3). With this approach, we deleted each inverted repeat element independently (ΔIR1 and ΔIR2), both inverted repeat elements (ΔIR1ΔIR2), and the A/T rich palindrome which centers the inverted repeat sequences (ΔPAL).

The modified DNA was assessed with Palindrome Analyzer for cruciform forming potential. This analysis revealed that each DNA construct still retained cruciform forming potential with comparable *ΔG* values ranging from 12.94 to 14.37 to the original cruciform (14.14) even with the nucleotide deletions (fig. S5). Proof of cruciform formation for these constructs was obtained by demonstrating T7 endonuclease cleavage of the cloned supercoiled DNA sequences (fig. 5a). This further substantiates the high propensity for A/T rich DNA (>90%) in intergenic regions to form cruciforms given appropriate base pairing and energetic provisions^24, 33^. Furthermore, the corresponding genetic deletions created different DNA juxtapositions for the *exsBA* intergenic DNA sequence which were predicted to alter HlyU binding. Indeed, purified HlyU was shown to bind inefficiently to the modified linear DNA fragments and required high amounts to initiate shifts in EMSA experiments (fig. 5b).

**Fig. 5.**
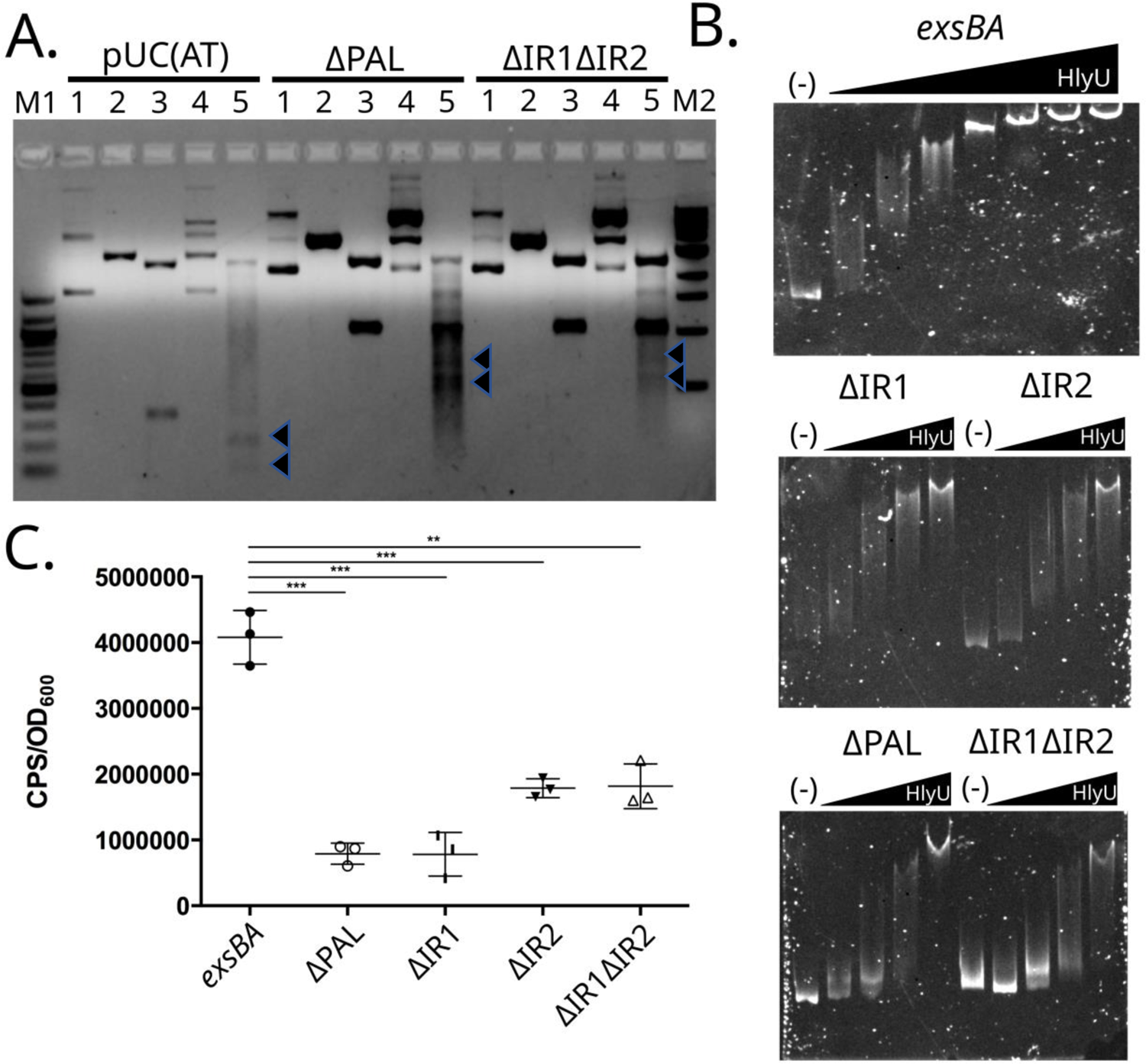
Genetic deletions of intergenic *exsBA* inverted repeat and palindrome sequences retain cruciform formation but are altered for HlyU binding efficiency. (A) Cruciform cleavage assays using T7 endonuclease and other restriction enzymes. 1-undigested plasmid, 2-linearized plasmid, 3-PvuII, 4-T7 Endonuclease, 5-T7 Endonuclease followed by PvuII. T7 Endonuclease targets cruciform structures to cause double-strand break in a two-step process. PvuII was used as it flanks the cloned DNA and allows for restriction mapping. Cruciform associated DNA fragments are indicated by arrowheads in lane 5 of the respective samples. M1 and M2 are DNA molecular size standards (B) EMSA with increasing amounts of purified HlyU protein mixed with the indicated *exsBA* genetic deletion fragments. The DNA species were stained with SYBR green. The (-) indicates no HlyU protein and therefore unshifted DNA template. (C) Luciferase activity of plasmid constructs with genetic transcriptional fusions to a *luxCDABE* cassette. The relevant *exsBA* intergenic deletions are indicated and ‘*exsBA*’ represents the wildtype sequence. Multiple t-tests were performed to determine statistical significance, ***: *p-value* < 0.01, **:*p-value*<0.05. All experiments were repeated at least three times. Technical replicates are shown in panel C.

Next, the corresponding modified *exsBA* intergenic DNA fragments were fused to a promoter-less *luxCDABE* cassette in a plasmid reporter system and assessed for luciferase expression in wild type *V. parahaemolyticus*. In this system, chromosomally encoded ExsA proteins act *in trans* to activate the cloned *exsA* promoter in the plasmid construct to support light emission. Moreover, HlyU proteins acting *in trans* interact with appropriate *exsBA* DNA and inform on cruciform attenuation. We did observe some HlyU-independent light emission from this plasmid system (i.e., within a Δ*hlyU* strain) however the level was significantly reduced compared to wild type *V. parahaemolyticus* (fig. S6) thus demonstrating that maximal gene expression was HlyU-dependent. All the modified *exsBA* intergenic DNA constructs were introduced in wildtype *V. parahaemolyticus* and were shown to produce reduced light emission compared to the wildtype *exsBA* intergenic sequence (fig. 5c). Notably, the ΔPAL construct, which forms a cruciform but is missing the 14 palindromic nucleotides that make up the central core of the HlyU binding site was the most negatively impacted for light expression. The ΔIR2 and ΔIR1ΔIR2 constructs supported intermediate levels of light emission, below that of wildtype *exsBA* DNA but higher than ΔPAL and ΔIR1. Combined with the T7 endonuclease cleavage results, these data indicate that different cruciforms produce repressive structures that impact on transcription activity. More importantly, the ability of HlyU to efficiently bind its target sequence has a positive effect on transcriptional activity, likely by competitively attenuating a cruciform structure.

### HlyU is required for ExsA auto-activation

The HlyU binding site partially overlaps the cruciform forming locus and constitutes a co-localized *cis*-genetic element that is involved in *exsA* gene expression. However, the location of HlyU binding is approximately 70 base pairs downstream of the known auto-regulatory *exsA* promoter^5, 9^. Based on this promoter positioning, it is unclear how HlyU positively impacts *exsA* promoter activity however alterations to local DNA topology induced by HlyU binding could be involved.

To better study this system and parse the role of HlyU that supports ExsA production, we reconstituted the *exsBA* intergenic region in a transcriptional luciferase reporter system within a heterologous *E. coli* strain (DH5αλ*pir*) which does not contain native HlyU or ExsA proteins. Recombinant plasmids coding for HlyU and ExsA were generated, and we confirmed the expression of HlyU and ExsA in this system by immunoblotting cell lysates (fig. 6a).

**Fig. 6.**
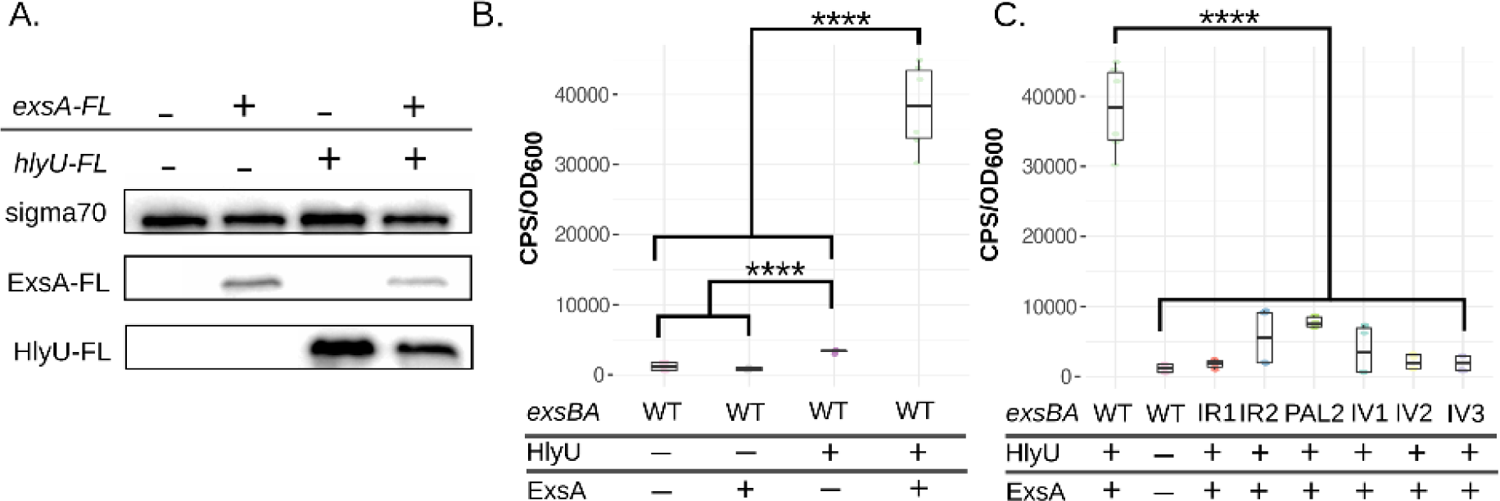
Reconstitution of the *exsBA* minimal regulon in *E. coli* provides evidence for a kick-start model of *exsA* regulation in *V. parahaemolyticus.* (A) *E. coli* DH5αλ*pir* cell lysates from strains harboring plasmids as indicated were subjected to immunoblotting to detect ExsA and HlyU. (B) DH5αλpir containing an *exsBA* transcriptional reporter fusion, *hlyU,* and/or *exsA* were assessed for light emission after 3 hours at 37°C. Photons per second (CPS) and cell density (OD_600_) were measured. (C) Bacterial strains containing cruciform mutations in context of the *exsBA-luxCDABE* transcriptional reporter fusion were assessed for light emission as in panel B. Multiple t-tests were performed to determine statistical significance, ****: *p-value* < 0.0001. Data was plotted with R and the ggstatsplot package.

As expected, expression of both HlyU and ExsA in *E. coli* supported robust transcriptional activity of the *exsBA* intergenic region as measured by the luciferase reporter (fig. 6b). By expressing HlyU alone in the heterologous system (without ExsA), we identified that HlyU was not only necessary, but also sufficient to drive low level luciferase expression (fig. 6b). This observation suggests the existence of an additional cryptic promoter downstream of the ExsA autoregulated promoter. Critically, ExsA alone was unable to generate luciferase expression from the *exsBA* intergenic region, revealing that HlyU is essential to support transcriptional activity driven from the ExsA autoregulated promoter. This suggested that HlyU binding to DNA is involved in the removal of a repressive DNA cruciform. Indeed, mutations that alter HlyU binding yet maintain cruciform forming potential were significantly reduced for *exsA-luxCDABE* expression (fig. 6c). These data suggest that HlyU binding is critical for cruciform attenuation which supports *exsA* expression. Moreover, these *E. coli* heterologous system data are in direct agreement with the inverted repeat deletion plasmid experiments performed in *V. parahaemolyticus* in figure 5.

### Transcriptional Start Site Mapping Identifies a Cryptic Promoter

The potential of a cryptic HlyU dependent promoter within the *exsBA* intergenic region was very interesting in that it would theoretically support initial ExsA production to autoregulate the ExsA-dependent *exsA* promoter (e.g. positive feedback loop). To address this possibility, we mapped transcriptional start sites (TSS) for mRNA species containing the *exsA* open reading frame using 5’-**<u>R</u>**apid **<u>A</u>**mplification of **<u>c</u>**DNA **<u>E</u>**nds (5’RACE). As expected, we identified *exsA* mRNA originating from near the known auto-regulatory distal *exsA* promoter (fig. S7a,b). Notably, we discovered a shorter *exsA* mRNA transcript initiating downstream of the HlyU binding site and cruciform forming locus (fig. S7a,b). We then confirmed the presence of a cryptic proximal *exsA* promoter (P_1_) using a luciferase reporter fusion. The entire *exsBA* intergenic region generated significant luciferase expression consistent with the existence of the distal auto-regulatory promoter and with previous experiments^9^ (fig. 7a). However, when the auto-regulatory distal promoter P_2_ was removed (ΔP_2_), the newly identified P_1_ promoter was still able to drive luciferase expression (fig. 7b). Importantly, a *hlyU* null mutant generated significantly less luciferase expression from the same luciferase reporter, indicating that the P_1_ promoter requires HlyU for maximal activity (fig. 7b). It is noteworthy that the HlyU-dependent P_1_ promoter is located within a DNA region that directly overlaps with H-NS binding (fig. S7a)^40^. Collectively, these data identify at least two distinct mRNA species which support *exsA* expression. These data also address how ExsA is first expressed through HlyU binding upstream of the P_1_ promoter, allowing ExsA to positively feedback and auto-regulate the distal promoter and therefore much of its own expression.

**Fig. 7.**
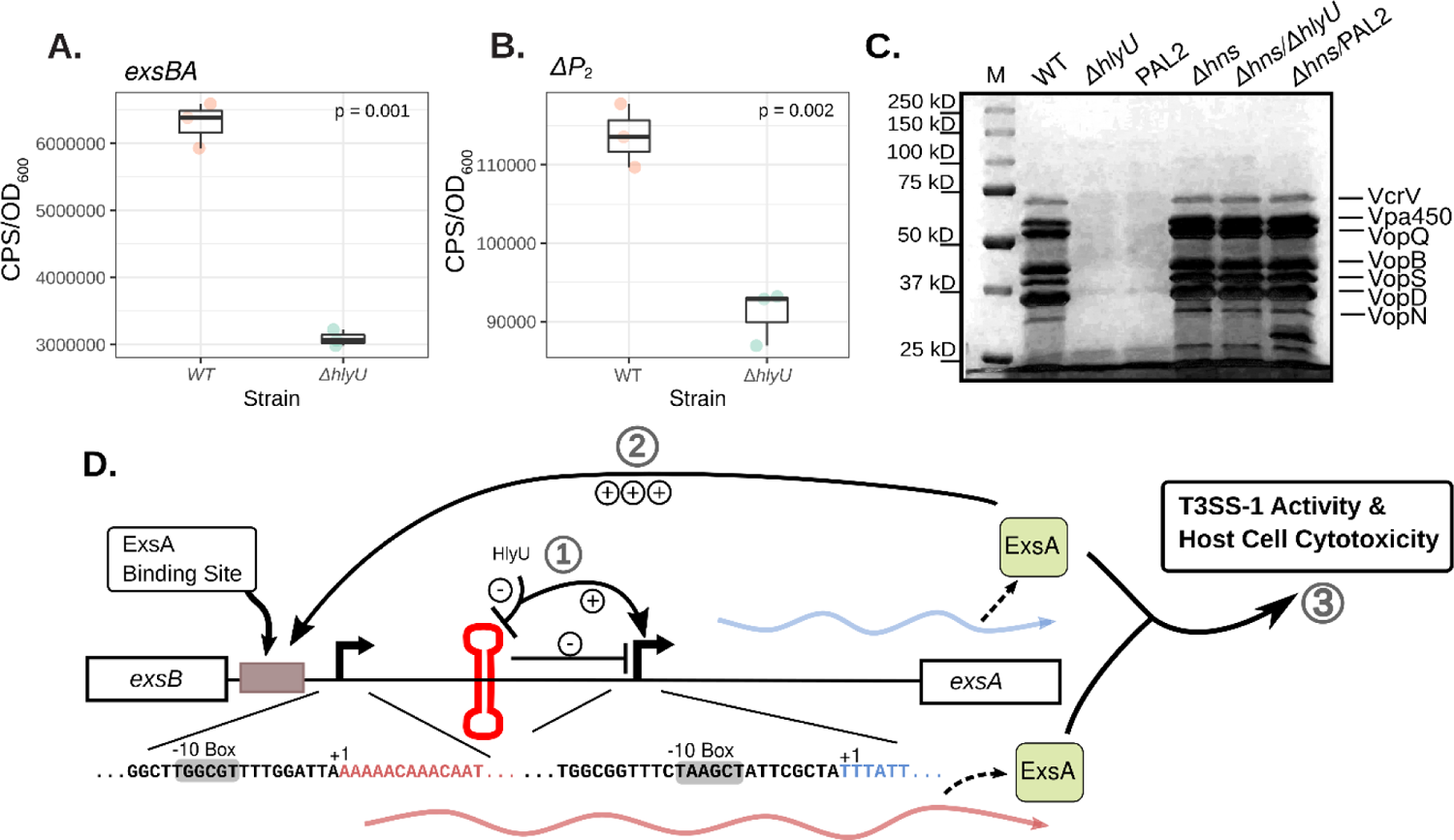
The *exsBA* intergenic region contains an autoregulatory and internal promoter elements that require HlyU binding near a cruciform to initiate a positive transcriptional feedback loop. (A) The *exsBA* intergenic region drives high level light emission from lux reporter construct in an HlyU dependent manner in *V. parahaemolyticus*. (B) Deletion of the autoregulatory promoter [P_2_] reveals light emission from an internal promoter that requires HlyU for maximal activity. (C) HlyU binding to a palindromic sequence near the cruciform locus is no longer required in the absence of H-NS to support T3SS-1 associated gene expression. Total secreted proteins of the indicated strains were subjected to SDS-PAGE and then stained by Coomassie blue. The indicated T3SS-1 proteins have previously been identified by mass spectrometry analyses. (D) Schematic model of genetic regulation of *exsA* within the *exsBA* intergenic region. 1) HlyU binds to DNA at the *exsBA* intergenic region and attenuates a cruciform structure allowing the activation of an internal promoter for *exsA* expression. 2) Expressed ExsA can autoactivate a distal upstream promoter, driving a majority of ExsA expression (3). Expressed ExsA can then drive T3SS-1 activity and host-cell cytotoxicity. DNA sequences shown indicate the identified promoter and transcriptional start sites from the described 5’RACE experiment.

### The action of HlyU binding at the cruciform locus is dispensable in the absence of H-NS

Interpretation of the overall genetic and biochemical data suggested to us that a transcriptionally repressive DNA cruciform within the *exsBA* intergenic region is attenuated by HlyU to support concomitant *exsA* gene expression. We and others have previously shown that *V. parahaemolyticus hns* mutants are deregulated for *exsA* expression and hyper-secrete T3SS-1 proteins ^9, 39^. H-NS has been shown to bind an approximately 200 bp span within the *exsBA* intergenic region^40^. It is possible that H-NS could mask the newly identified P1 promoter from cellular transcriptional machinery (fig. S7a). The exact mechanism of H-NS gene silencing of *exsA* expression is unknown but based on the large section of DNA coverage, it likely involves H-NS nucleation that leads to nucleofilament mediated DNA ‘stiffening’ to silence transcription^50^. Moreover, H-NS contributes to constraining chromosomal DNA supercoiling^23^ which could also impact on *exsA* expression. In support of this view, magnesium, which is known to reduce H-NS associated nucleofilaments ^30, 51^ is a potent inducer of *exsA* expression and T3SS-1 activity in wild type *V. parahaemolyticus*^49^. Nonetheless, any influence of magnesium on H-NS must be synergistic with contextual HlyU action to support T3SS-1 activity as *hlyU* null mutants are defective for *exsA* expression and efficient T3SS-1 secretion^9^ (see fig. 4a). We attempted to rescue a *hlyU* mutant for T3SS-1 activity by culturing it in elevated amounts of magnesium (up to 55mM) which inhibits H-NS binding to DNA *in vitro*^51^ but were not successful in restoring *exsA* expression (data not shown). This further highlighted the critical role of HlyU for *exsA* expression and suggested synergistic actions of HlyU binding to DNA along with a separate undefined H-NS de-repression mechanism for efficient T3SS-1 expression.

We set out to investigate the T3SS-1 associated regulatory effect of HlyU binding to DNA in the presence and absence of H-NS. Notably, H-NS contributes to constraining DNA supercoiling which is a requisite condition for DNA cruciform formation^37^. We hypothesized that the important role of HlyU binding to *exsBA* intergenic DNA would be dispensable for T3SS-1 activity in the absence of H-NS. Indeed, a Δ*hns*Δ*hlyU* strain exhibited deregulated T3SS-1 activity revealed by elevated T3SS-1 proteins in a secretion assay (fig. 7c). Further, in a Δ*hns/*PAL2 double mutant where HlyU is present but unable to efficiently bind *exsBA* intergenic DNA due to alteration of the target palindrome sequence, similarly exhibited high levels of T3SS-1 secreted proteins (fig. 7c). In stark contrast, the same PAL2 mutation in the presence of HlyU and H-NS was unable to support efficient T3SS-1 protein secretion (i.e. PAL2, fig. 7c), revealing a context specific H-NS associated phenotype for this mutation. This suggests that HlyU binding near the site of cruciform formation is a critical event for productive *exsA* expression in wild type *V. parahaemolyticus* with a condensed nucleoid under typical torsional stresses. In the absence of H-NS, the strict requirement for HlyU binding to *exsBA* intergenic DNA is eliminated. This suggests that in *V. parahaemolyticus*, a separate but synergistic mechanism of HlyU-mediated cruciform attenuation and H-NS activity contributes to *exsA* gene expression and T3SS-1 associated virulence.

## Discussion

This study reports HlyU DNA-binding at the site of a DNA cruciform as the first step in an extensive regulatory cascade leading to T3SS-1 virulence gene expression in *Vibrio parahaemolyticus*. We propose the following mechanism for DNA cruciform involvement in *exsA* expression and T3SS-1 activity in *Vibrio parahaemolyticus:* 1) HlyU binding to DNA attenuates a repressive DNA cruciform which supports internal promoter activity leading to initial low level *exsA* mRNA expression, 2) initial ExsA production (mediated by HlyU function) autoregulates the upstream *exsA* promoter to drive elevated *exsA* transcription, and 3) high cellular levels of ExsA drives the expression of multiple ExsA-dependent T3SS-1 gene operons^5^ leading to host cell cytotoxicity (fig. 7d).

In many pathogenic *Vibrio* species, HlyU has been proposed to alleviate H-NS mediated virulence gene silencing by outcompeting and displacing H-NS from A/T rich DNA sequences^17, 18^. This view agrees with reports for *Vibrio* spp. DNA binding regulators such as LuxR and ToxT^19, 52^ and aligns with the roles of DNA binding regulators in other pathogens (e.g., SsrB, Ler^53, 54^). It is reasonable to consider that each DNA binding regulator displaces H-NS associated DNA in a contextual and localized manner. Moreover, H-NS nucleofilament DNA formation at A/T rich regions and DNA bridging mechanisms introduce negative DNA supercoiling and localized DNA structural changes^30^. Such conditions overcome the energetics of DNA duplex stability to generate unpaired DNA bases such as those found in cruciforms. We recognize our data for DNA cruciform formation always depended on DNA supercoiling whether it was *in vitro* (purified supercoiled plasmid DNA) or *in vivo* (CAA treated bacteria). Critically, these observations agree with previous DNA supercoiling mechanistic studies and A/T rich sequence requirements for DNA cruciform formation^24, 37, 44^.

DNA cruciforms are very challenging to study due to the localized energy input and base pairing requirements that contribute to their formation. We used a variety of traditional genetic and biochemical tools in addition to pharmacological (novobiocin) and chemical genetic approaches (chloroacetaldehyde treatment) to thoroughly investigate the DNA cruciform at the *V. parahaemolyticus exsBA* locus. Our T7 endonuclease cruciform cleavage data was in striking agreement with model oligonucleotide J-structure cruciform cleavage studies^46^. Specifically, we observed multiple T7 endonuclease cruciform cleavages 3-5 nucleotides from the cruciform base (fig. 1c), which is the established sequence independent location of cleavage for this enzyme. Additional independent evidence for single stranded ‘loop’ DNA associated with cruciforms was obtained with mung bean nuclease digestions. We also found T7 endonuclease cleavage evidence for cruciform forming DNA in proximity to HlyU binding sites in *V. cholerae* (*hlyA)* and *V. vulnificus* (*rtxA1* operon region), but not for *V. anguillarum* (*rtxH-rtxB)*. Palindrome analyser indicated the putative *V. anguillarum* cruciform as requiring more energy input for cruciform formation (*ΔG*=17.29) than that seen in the other *Vibrio* spp. (*ΔG*=14.14 for *V. parahaemolyticus,* fig. 1c). This provides two possibilities: 1) that a cruciform structure simply doesn’t exist at this site in *Vibrio anguillarum* or 2) that our plasmid system is incapable of creating the necessary conditions for cruciform formation outside of the *V. anguillarum* cell. Nonetheless, DNA cruciforms are found near HlyU binding sites in multiple pathogenic *Vibrio* spp. In the case of *V. parahaemolyticus exsBA* genetic locus, the DNA cruciform appears to operate as a transcriptionally repressive element that requires attenuation to permit *exsA* gene expression from a cryptic internal promoter that is masked by H-NS. As T3SS-1 biosynthesis is a major cellular investment in *V. parahaemolyticus* (expression of 40+ genes)^5^, it follows that the entry master regulator ExsA is tightly repressed to prevent spurious expression. Furthermore, *exsA* expression is contextually de-repressed by HlyU to support coordinated T3SS-1 associated gene expression during host infection.

Our data expands knowledge relating to HlyU-DNA binding outcomes. The DNA binding data along with HlyU crystal structures suggest that a core A/T rich palindrome forms DNA major grooves to accommodate α4 helical domains found within wHTH HlyU dimers^14, 38^. Furthermore, inverted repeat DNA elements are implicated in binding the HlyU ‘wing’ domains to support efficient binding^12^. Notably, the IR1 mutant *V. parahaemolyticus* strain presented in this study is particularly interesting as it produced an intermediate level of T3SS-1 activity and host cytotoxicity compared to wild type bacteria. The IR1 mutation alters the inverted DNA repeat sequence on the left side of central core palindrome, while the right inverted DNA repeat element IR2 is unchanged (fig. 3b). Our data indicates that HlyU achieves incomplete and imperfect binding in context of IR1 altered DNA (fig. 3c). We speculate that this imperfect binding likely supported partial attenuation of cruciform formation, thus allowing for partial *exsA* expression. Critically, a similar outcome was not observed for the alterations of the DNA inverted repeat to the right of the core palindrome DNA (i.e., IR2 mutants). The key difference between IR1 and IR2 DNA elements is that IR1 is outside of the cruciform forming DNA stem and loop sequences, whereas IR2 directly overlaps and is within the cruciform DNA locus (fig. 1d). While additional studies will be required to unravel these experimental observations, it appears that HlyU binding in the immediate vicinity of the cruciform locus produces DNA topology changes that contextually reveal a cryptic promoter leading to *exsA* gene expression.

Our data supports two models for *V. parahaemolyticus* HlyU interaction with cruciform forming DNA. In the first model, perhaps the simplest, HlyU interacts with double-stranded DNA and as such prevents the formation of a DNA cruciform by sterically hindering annealing of cruciform stem associated nucleotides. This notion is best supported by our data, most notably by DNA-binding EMSA analysis using linear dsDNA. The requirement of HlyU for *exsA* promoter activation *in situ* (fig. 6b) is also consistent with this interpretation. An alternative more complex possibility is that HlyU interacts with bent DNA found at the base of the cruciform structure and destabilizes the cruciform directly by conformational changes induced by protein binding. Such an interaction is possible based on studies evaluating the formation and structure of DNA cruciforms^24, 33, 37, 46^ and HlyU binding to a bent planar face of DNA^12, 14^. We explored this possibility with *in vitro* studies but were unsuccessful in showing a direct HlyU interaction with a synthetic cruciform DNA structure (data not shown). We cannot exclude the possibility that the synthetic DNA cruciform was improperly formed or assumed a conformation less favorable for HlyU binding. Therefore, the presented data has limitations in that it does not identify a defined HlyU mechanism for direct cruciform binding. Rather the data suggests that cruciform attenuation is necessary for initial *exsA* gene expression and the process is strictly dependent on HlyU binding to DNA. Complex biophysical protein-DNA binding experiments beyond the scope of this study will be required to test the proposed models in future studies.

DNA cruciforms and other 4-way junctions are found in all living cells and some plasmids and viruses. The dynamic and temporal aspects of cruciform formation are modestly understood however DNA supercoiling is thought to provide energy to facilitate DNA strand extrusion at A/T rich containing regions^37^. This outcome would likely occlude or prevent RNA polymerase access to specific DNA promoters^24^. Accordingly, specialized DNA binding proteins would be required to act near certain DNA cruciforms to permit transcription related activities. The data presented here newly implicate HlyU in destabilizing a transcriptionally repressive DNA cruciform thus contextually supporting access to a previously silenced genetic promoter. Considering that chromosomal DNA supercoiling and A/T rich DNA are common features of intergenic regions^32, 34, 37, 41, 42^, we believe that cruciform attenuation, driven by specialized DNA binding proteins, may represent an overlooked extricating mechanism to de-repress gene expression.

## Materials and Methods

### Bacterial cultures and growth conditions

*Vibrio parahaemolyticus* RIMD2210633 was cultured in either LB-Miller (10g/L tryptone, 5g/L yeast extract, 10g/L NaCl) or LBS (10g/L tryptone, 5g/L yeast extract, 20g/L NaCl, 20mM Tris-HCl, pH 8.0). Antibiotic concentrations used for *V. parahaemolyticus* are as follows: chloramphenicol (2.5 µg/mL). *V. parahaemolyticus* was cultured at room temperature (∼22°C), 30°C, or 37°C, depending on a given experiment’s requirements. *E. coli* strains were cultured in LB-Miller at 37°C containing the following antibiotics when necessary: ampicillin (100 µg/mL), chloramphenicol (30 µg/mL), and tetracycline (5 µg/mL).

### Recombinant DNA approaches

PCR and DNA cloning was performed using standard techniques. All DNA polymerases and restriction enzymes were purchased from New England Biolabs (NEB) unless stated otherwise. Control cloning and restriction digestion experiments were performed in parallel for interpretation purposes.

### *In silico* Cruciform Analysis

Intergenic sequences known to contain HlyU binding sites by previous studies were selected from various *Vibrio* spp. and were used as input into Palindrome Analyser - an online bioinformatics tool which identifies cruciform forming DNA sequences and calculates the amount of energy required for cruciform formation^43^. Our cruciform identification required a cruciform stem of at least 6 base pairs, with a spacer/loop region of at least 10 bp. We allowed up to a single mismatch in the cruciform stem sequences. For each sequence, the 10 (or fewer) possible cruciforms requiring the smallest change in free energy to form are detailed.

### T7 Endonuclease and cruciform restriction mapping assays

HlyU binding sites within *Vibrio sp*. intergenic DNA regions were evaluated for cruciform structures by cloning PCR amplified DNA fragments or synthetic gBlocks (Integrated DNA Technologies (IDT), see Table 3) into pBluescript and transforming the recombinant plasmid DNA into *E. coli* DH5α. Freshly prepared supercoiled plasmid DNA was then isolated from overnight cultures using a Monarch miniprep kit (NEB) and then immediately subjected to restriction digestion with T7 endonuclease (NEB) which cleaves at DNA cruciforms. To determine the approximate localization of DNA cruciform structures, a sequential digest with PvuII (following an initial T7 endonuclease digestion) was performed as previously described^45^ followed by agarose gel electrophoresis to resolve digested DNA fragments.

To precisely detect cruciform cleavage sites, we designed a strategy using T7 endonuclease digestion or mung bean nuclease, the latter which digests single stranded ‘loop’ DNA within DNA cruciforms^35^. Briefly, supercoiled plasmid DNA was treated with T7 endonuclease followed by reaction purification and treatment with mung bean nuclease to remove ssDNA overhangs and blunt the DNA to a T7 endonuclease cleavage site. Linearized dsDNA was then selectively extracted from an agarose gel, subjected to PvuII digestion (2 PvuII sites flank the DNA cruciform) yielding two blunt ended DNA fragments which were separately cloned into EcoRV treated pBluescript. A similar approach using only mung bean nuclease to detect ssDNA within cruciform ‘loops’ was also pursued. DNA sequencing of the resultant recombinant plasmids revealed DNA cruciform cleavage sites for either T7 endonuclease or mung bean nuclease as indicated by ligation to the EcoRV pBluescript site.

### Transcriptional reporter assays using exsBA DNA with altered HlyU binding potential

Synthetic gene fragments (IDT) were designed with specific nucleotide deletions that were predicted to impact HlyU binding (Table#). The HlyU binding site is composed of two perfect inverted repeats (ATATTAG, CTAATAT) that flank a central (core) A/T rich palindromic sequence (TAATTTAATTTATT). Each of the altered *exsBA* DNA fragments along with a wild type *exsBA* fragment were separately cloned into a plasmid containing a promoter-less *luxCDABE* cassette to create transcriptional reporter constructs. The corresponding plasmids were incorporated into wild type *V. parahaemolyticus*. The bacteria were then cultured under T3SS-1 inducing conditions (magnesium supplementation and EGTA)^49^ to induce *exsA* expression. Luciferase activity derived from the lux cassette was measured as light emission 2.5 hours post induction using a Victor X5 luminometer as previously described^55^.

### *Vibrio parahaemolyticus* chromosomal mutant generation

Δ*hlyU* and Δ*hns* null chromosomal mutants were generated using allelic exchange and sucrose selection as previously described^9^. Multiple DNA fragments (IDT, synthesized gBlocks) with specific nucleotide substitutions were designed to mutate HlyU binding site or cruciform forming locus within the *V. parahaemolyticus exsBA* intergenic region on the chromosome (see table S5, table S7 and fig. 3b). The DNA fragments were cloned into suicide plasmid pRE112 and used in allelic exchange experiments. All derived mutant strains were verified by DNA sequencing of PCR amplified chromosomal DNA to confirm genetic changes.

### *in situ* chromosomal cruciform modification by chloroacetylaldehyde (CAA) pulse-chase treatment

Cell permeable CAA was used to chemically modify *in situ V. parahaemolyticus* chromosomal DNA cruciforms by generating nucleotide base ethenoadducts at specific sites^48^. Specifically, a CAA treatment (pulse) reacts with unpaired bases within DNA cruciforms resulting in localized DNA damage. Upon removal of CAA (chase) the damaged DNA can be assessed for effects on locus-specific gene expression. In this pulse-chase approach, 1 x10^9^ stationary phase cells from an overnight culture were harvested by centrifugation, washed in PBS (Phosphate Buffered Saline), and then treated with 40nm CAA (or mock-treated with PBS) for 30 minutes. The cells were washed twice with PBS to remove CAA, and then immediately used to measure *exsA* promoter activity by an *in situ* real-time quantitative luciferase reporter assay. Cells within the population with CAA damaged DNA cruciform lesions were expected to emit less light than a paired mock-treated sample. We independently confirmed that a 30-minute 40nm CAA pulse treatment did not impair population-based cell growth metrics as determined by OD_600_ readings whereas higher concentrations were inhibitory and not pursued further (data not shown). Moreover, maintenance of CAA treatment during active growth conditions was not possible due to unpaired nucleotide bases associated with DNA replication events and various RNA species. Importantly, the assay is population based so cells that grow and repair CAA damaged DNA or suffer mutations are accounted for in the captured data.

### *In vitro V. parahaemolyticus* T3SS-1 protein secretion assay

T3SS-1 protein secretion assays were performed as previously described^49^. A characterized T3SS-1 defective strain Δ*vscN1* along with Δ*hlyU*^9^ served as relevant controls allowing for secreted protein profile comparisons.

### *V. parahaemolyticus* host cell cytotoxicity assays

HeLa cells (ATCC) were seeded in a 24-well dish at a density of 100,000 cells/mL and cultured for 16 hours. Overnight cultures of *V. parahaemolyticus* strains were grown in LB-Miller at 37°C and diluted in DMEM (Dulbecco Modified Eagle’s Medium, Invitrogen #11995) to generate a multiplicity of infection (MOI) of ∼2. The cultured HeLa cells were rinsed twice with phosphate-buffered saline (pH 7.4, ThermoFisher; 10010023), followed by the addition of the relevant bacterial cells to initiate infection. The infection was incubated for 4 hours at 37°C/5% CO_2_. Infection supernatants were collected and subjected to centrifugation (15000xg for 1 minute) to remove bacteria and HeLa cells. The fluorescent CyQUANT™ LDH Cytotoxicity Assay (ThermoFisher; C20302) was used to measure released lactate dehydrogenase (LDH) within the clarified supernatants as directed by the manufacturer. Percent cytotoxicity was calculated according to the following formula:

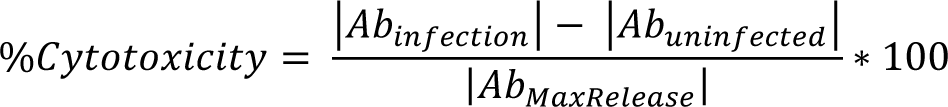

### Recombinant HlyU-His Protein Purification

*E. coli* BL21(λDE3) containing a cloned *V. parahaemolyticus hlyU-his* plasmid DNA construct (*8*) was cultured overnight. The following day, the bacteria were sub-cultured (1/50) and grown to an OD_600_ of 0.8 and then induced with 0.4mM IPTG and cultured for an additional 3 hours. The bacteria were harvested by centrifugation and the cell pellet was frozen at −20°C. Cell lysates were prepared, and nickel affinity chromatography was performed under soluble conditions as previously described^9^. Purified HlyU-His was subjected to Amicon ultrafiltration to enrich for dimeric HlyU (∼22 kDa). Protein expression and purification steps were assessed by SDS-PAGE analysis (SFig 5).

### Electrophoretic Mobility Shift Assays

Electrophoretic mobility shift assays were performed as previously described^9^. Briefly, DNA oligonucleotides (sequences found in Table 1) were mixed at equimolar concentrations and NaCl was added to a concentration of 50 mM to promote proper annealing of the oligonucleotides. The oligonucleotides were heated to 98°C followed by sequential cooling (1°C/5s) to 10°C thus generating annealed short dsDNA fragments.

A DNA master-mix in 1X EMSA buffer (1 mM Tris, 6 mM NaCl, 0.5 mM MgCl2, 0.01 mM EDTA, 0.1 mM CaCl2, 0.2% glycerol) was created for each dsDNA fragment and mixed with variable HlyU-His protein amounts or buffer alone to a final volume of 15µL. Reactions were allowed to equilibrate at room temperature for 30 minutes. Each reaction was subjected to TBE-PAGE for 1 hour at 100V at 4°C. Gels were stained with 1X SYBR Green (Invitrogen) for 30 minutes, rinsed in distilled H_2_O, and visualized with the BioRad VersaDoc platform.

### 5’ RACE

To determine mRNA transcriptional start sites in the *exsBA* intergenic region, a 5’ RACE experiment was performed as previously described^56^. *Vibrio parahaemolyticus* RIMD2210633 was inoculated at a starting OD of 0.025 in Mg/EGTA containing LB and incubated at 30⁰C/250RPM for 3 hours, prior to the isolation of total RNA. Total RNA was used for reverse transcription and PCR amplification. Primer AL400 (an *exsA* specific primer targeting the 3’ end) was used for reverse transcription, and **<u>G</u>**ene **<u>S</u>**pecific **<u>P</u>**rimers (GSP) 1 and 2 were designed to target the *exsA* coding region of mRNA templates (table S3).

### Reconstitution of *exsBA* genetic locus transcriptional activity in *E. coli*

Luciferase based reporter plasmids containing modified *exsBA* sequences were generated using standard cloning techniques (table S5) and were based on a verified *exsBA-luxCDABE-* VSV105 transcriptional reporter plasmid^9^. *pHlyU-*FLAG which expresses C-terminal FLAG epitope tagged HlyU from a recombinant *tac* promoter was transformed into *E. coli* DH5αλpir. To generate a FLAG epitope tagged ExsA expression construct driven by the *lac* promoter, the *exsA* gene was PCR amplified from *V. parahaemolyticus* chromosomal DNA using oligonucleotides NT387 and NT388 followed by cloning into pFLAG-CTC. This plasmid served as template DNA in a PCR with primers NT472 and NT473 to generate *exsAFL-*pRK415 (for ExsA-FLAG expression). All plasmid constructs were verified by DNA sequencing. The expression plasmids or empty control plasmids were transformed into DH5αλpir in various combinations together with appropriate *exsBA* intergenic DNA luciferase reporter plasmids (table S2).

Luciferase assays were performed by inoculating fresh LB with OD_600_ normalized overnight *E. coli* cultures allowing cell growth to mid-log phase (∼3 hours). 100 uL of culture was used to measure light emission measured as counts per second along with cell density readings (OD_600_). Each culture was measured in triplicate and the experiment was repeated twice to attain statistical significance by multiple t-test. Graphs were plotted using R and the ggstatsplot package. An immunoblot was used to detect FLAG epitope tagged HlyU and ExsA from cell lysates. Anti-FLAG (Sigma) and anti-sigma70 (BioLegend) antibodies were used as primary antibodies, and goat anti-mouse HRP (Cell Signaling Technology) as secondary antibodies. Images were captured using a Bio-Rad VersaDoc system.

## Acknowledgments

The authors would like to thank Ken Jarrell, Craig McCormick, John Rohde, John Archibald, and Andrew Roger for their insightful comments on the initial drafts of our manuscript. LJG is funded by a Vanier Canadian Graduate Scholarship, and a Killam PreDoctoral scholarship. LS and AC were funded by the National Science and Engineering Research Council (NSERC) Undergraduate Summer Research Award program. NAT holds an NSERC Discovery Grant-RGPIN 05807. Conceptualization: LJG, JMB, NAT Methodology: LJG, NAT Validation: LJG, NAT Formal analysis: LJG, LS, JMB, JM, AC, NAT Investigation: LJG, JMB, LS, AC, JM, NAT Writing – original draft preparation: LJG, NAT Writing – review and editing: LJG, NAT Visualization: LJG, LS Supervision: NAT, LJG

## Supplementary Materials

**Fig. S1.**
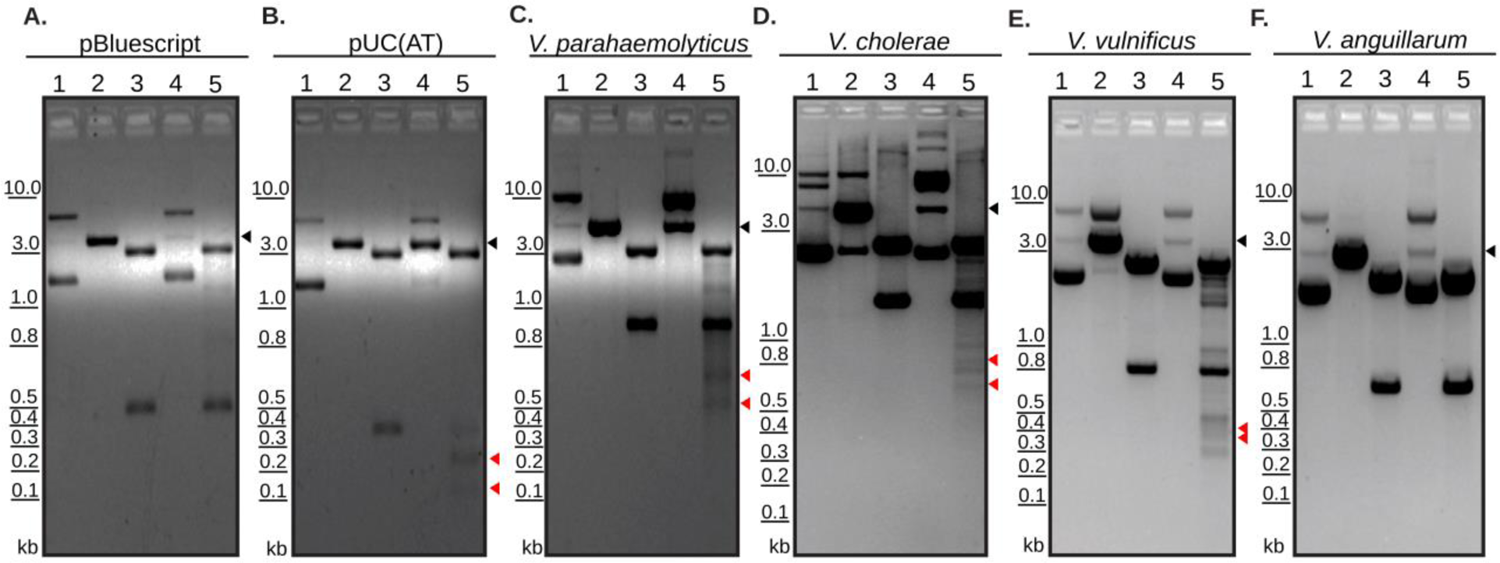
Identification of cruciform structures at intergenic regions in a variety of *Vibrio* spp. Restriction digestion of cloned DNA from *Vibrio* spp. visualized by agarose gel electrophoresis. 1-undigested plasmid, 2-linearized plasmid, 3-PvuII, 4-T7 Endonuclease, 5-T7 Endonuclease followed by PvuII. T7 Endonuclease targets cruciform structures to cause double-strand break in a two-step process. PvuII was used as it flanks the cloned DNA using sequences found in the plasmid backbone and allows for restriction mapping. Red arrowheads in lane 5 of each panel indicate DNA fragments released by digestion by PvuII and T7 endonuclease. Black arrowheads indicate the linearized DNA fragment as shown in lane 2. pBluescript (**A**) and pUC(AT) (**B**) are negative and positive controls respectively. Other cloned DNA from *Vibrio* spp. Includes *Vibrio parahaemolyticus exsBA* (**C**), *V. cholerae tlh-hlyA* (**D**), *V. vulnificus rtxA1* operon region (**E**), *and V. anguillarum plp-vah* (**F**).

**Fig. S2.**
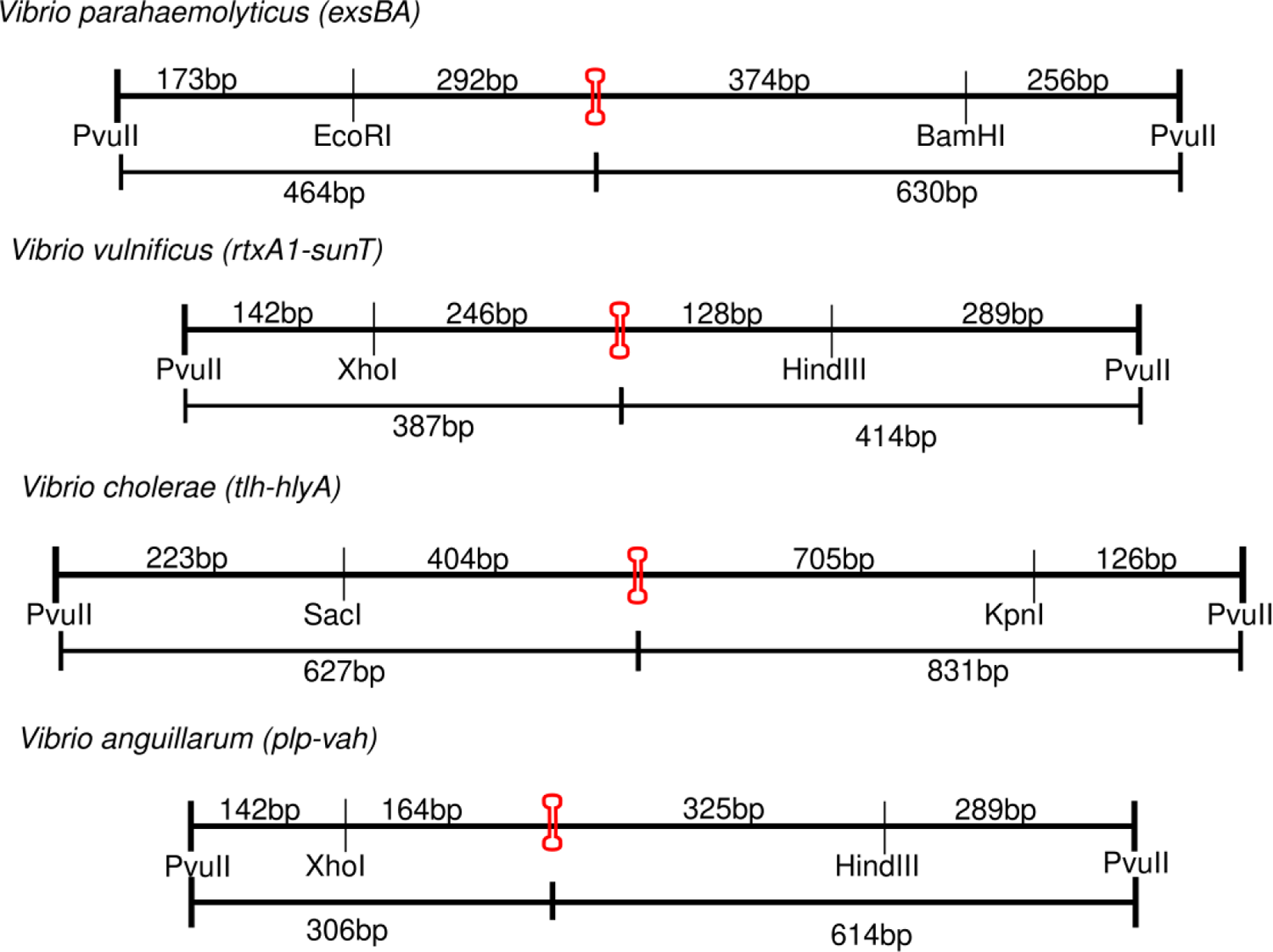
Restriction maps of cloned *Vibrio* spp. DNA fragments. The terminal PvuII sites are found within pBluescript, whereas the denoted internal restriction sites were used for cloning *Vibrio* spp. DNA into the vector’s multiple cloning site. The lowest free energy cruciform for each cloned DNA fragment is shown in red. In the case of *V. anguillarum*, although cruciform structures were identified in silico, cruciform cleavage was not detected (Fig S1).

**Fig. S3.**
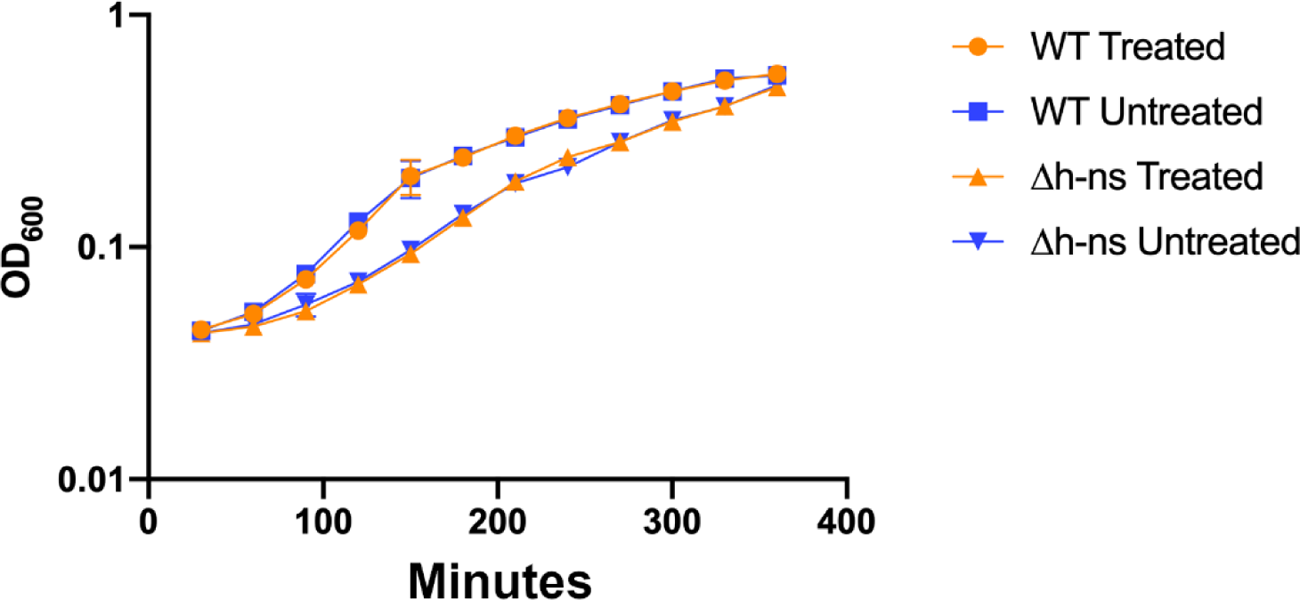
Growth curve for specified *V. parahaemolyticus* strains used within the chloroacetylaldehyde pulse-chase experiment. *exsBA-lux* strains (WT) or *exsBA-lux* (Δ*hns*) were treated with chloroacetylaldehyde (CAA) or PBS (negative control, untreated). No statistical difference was apparent between treated and untreated strain growth rates over a 380-minute period.

**Fig. S4.**
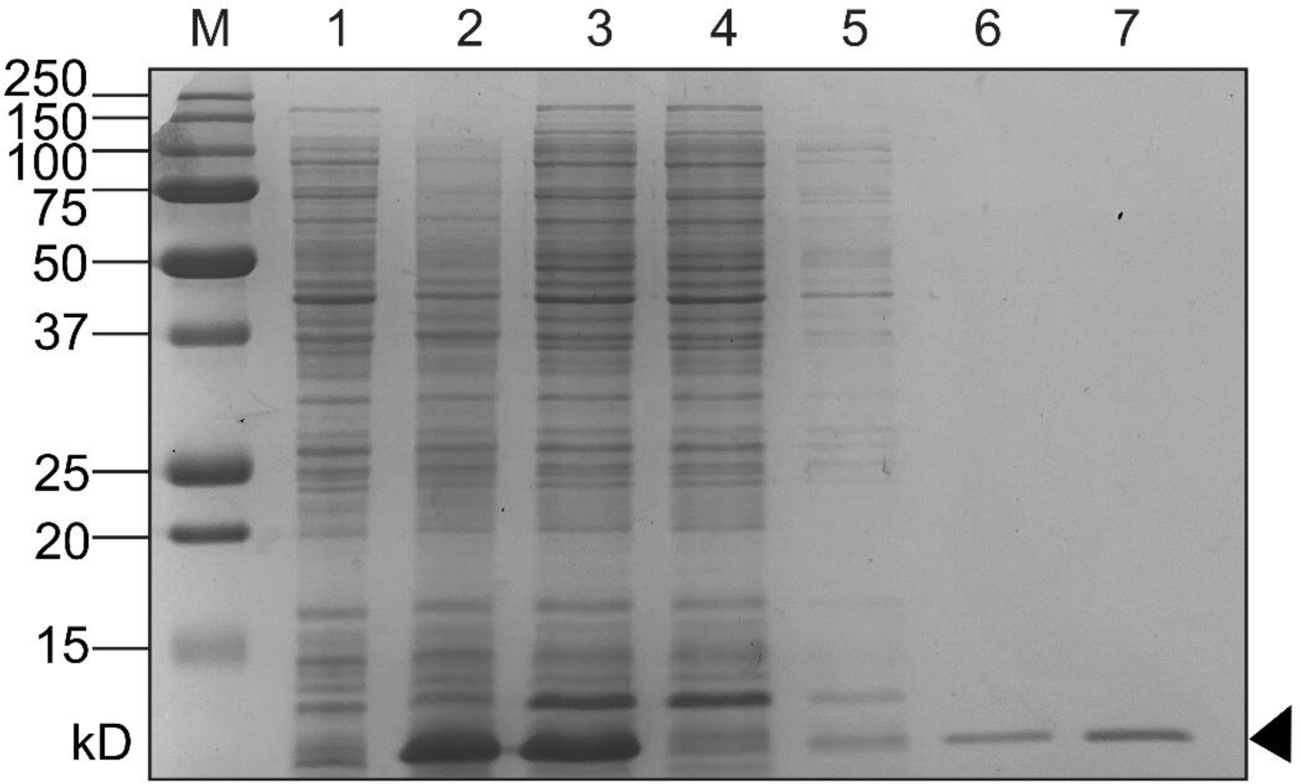
HlyU-His Protein Purification. Nickel affinity chromatography was used to purify His-tagged HlyU expressed using an IPTG induction protein expression system in BL21(λDE3) *E. coli*. Coomassie stained SDS-PAGE lanes are as follows: M – All Blue Protein Marker (Biorad), 1 – pre-induction cell lysate, 2 – Post-IPTG induction cell lysate, 3 – pre-column soluble lysate, 4 – post-column flowthrough, 5-Wash fraction (i), 6 – Wash fraction (ii), 7 – Elution fraction containing purified HlyU-His.

**Fig. S5.**
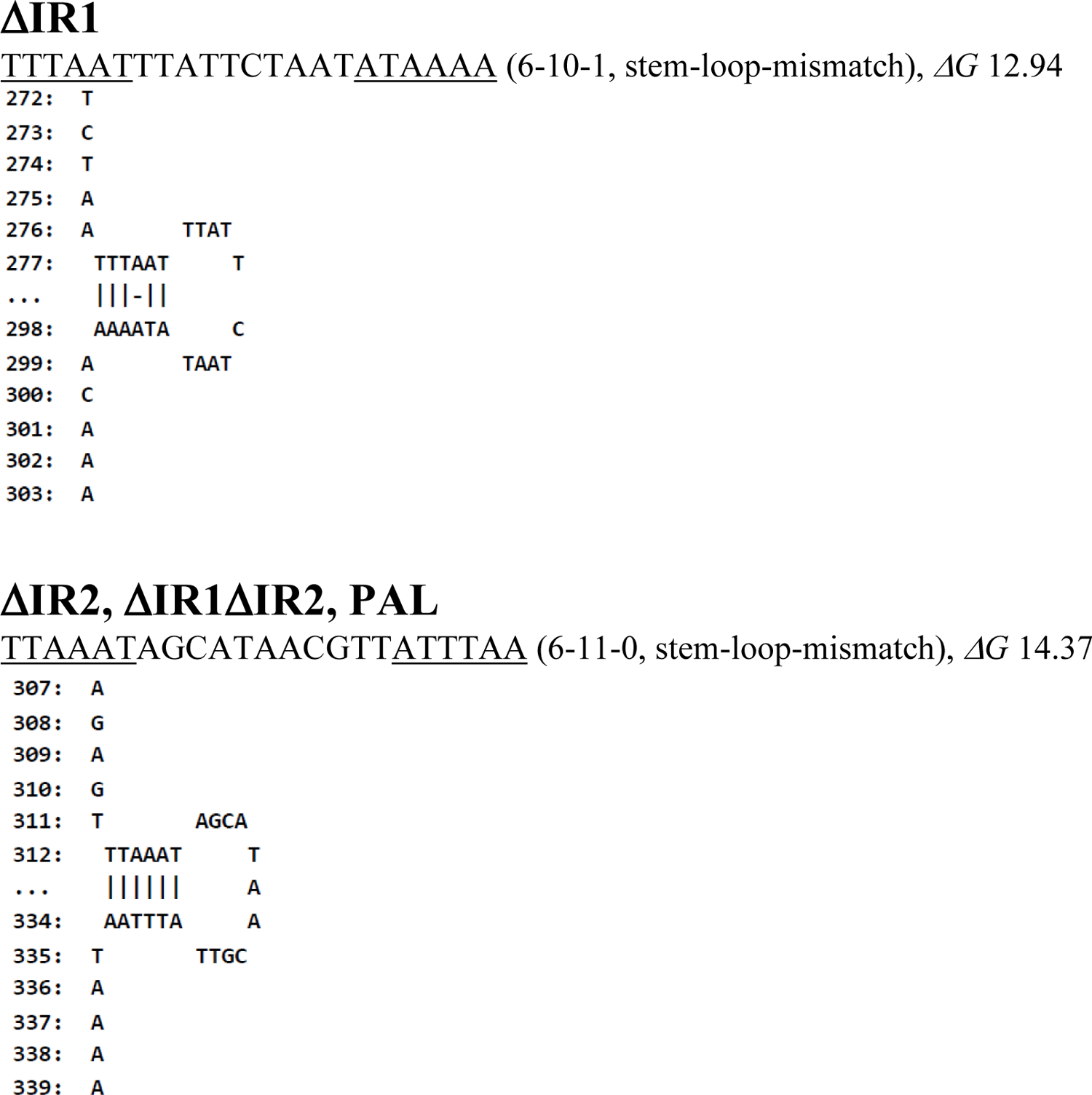
Palindrome analyzer results for *exsBA* inverted repeats and central palindrome genetic deletions. The lowest energy cruciforms are shown. The deletion of the central palindrome alone, or inverted repeat 2 (IR2) resulted in a DNA juxtaposition that created a cruciform forming element (6-11-0) for three *exsBA* constructs (ΔIR2, ΔIR1ΔIR2, and PAL).

**Fig. S6.**
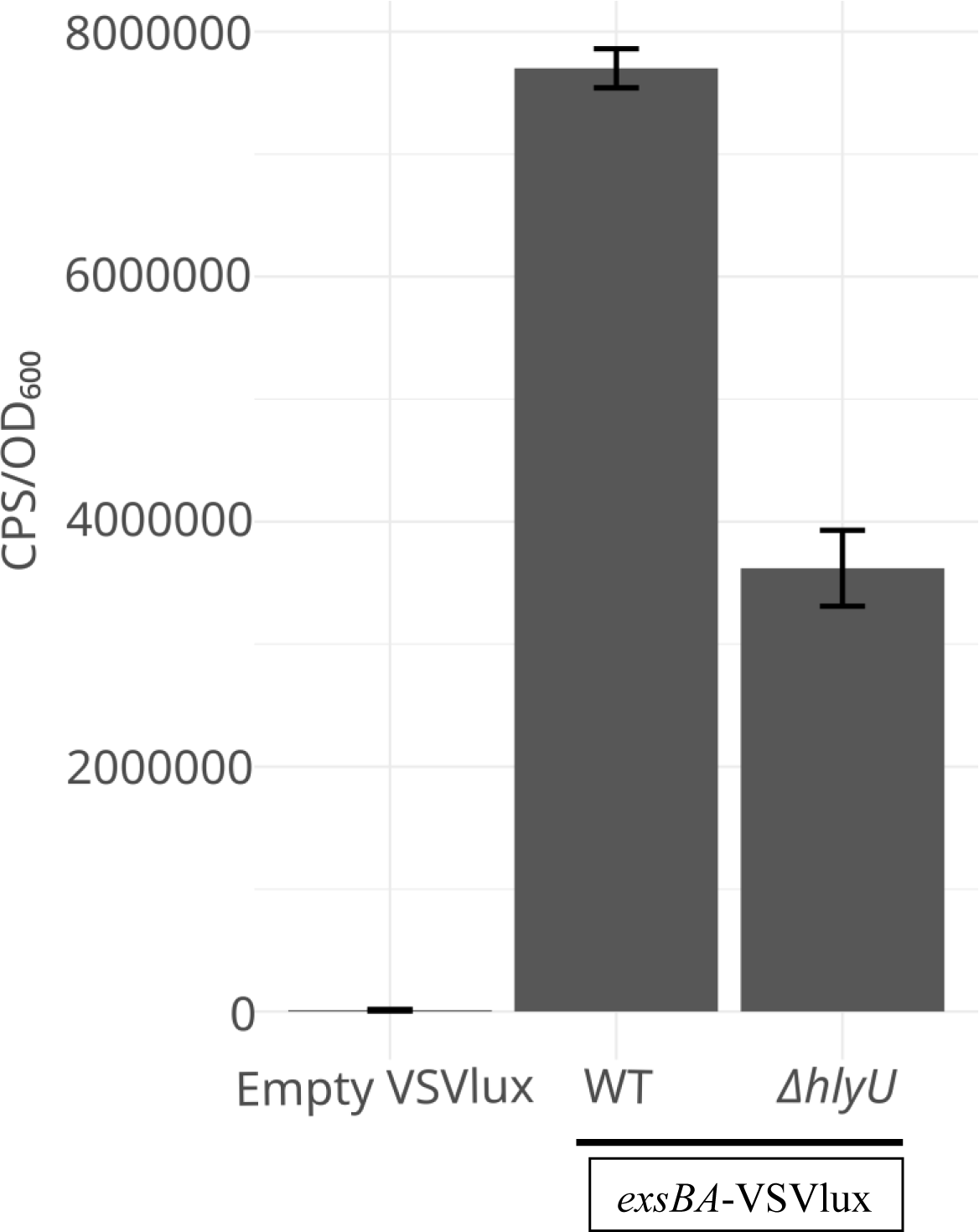
A plasmid-based transcriptional fusion of the *exsBA* intergenic region to a *luxCDABE* cassette is dependent on HlyU for maximal activity in *V. parahaemolyticus.* Some background activity occurs in the absence of HlyU presumably due to plasmid DNA replication, and changes in supercoiling during bacterial growth. The measurements were taken 2.5 hours post induction (magnesium +EGTA) which corresponds to the maximal activity observed for wildtype (WT) bacteria.

**Fig. S7.**
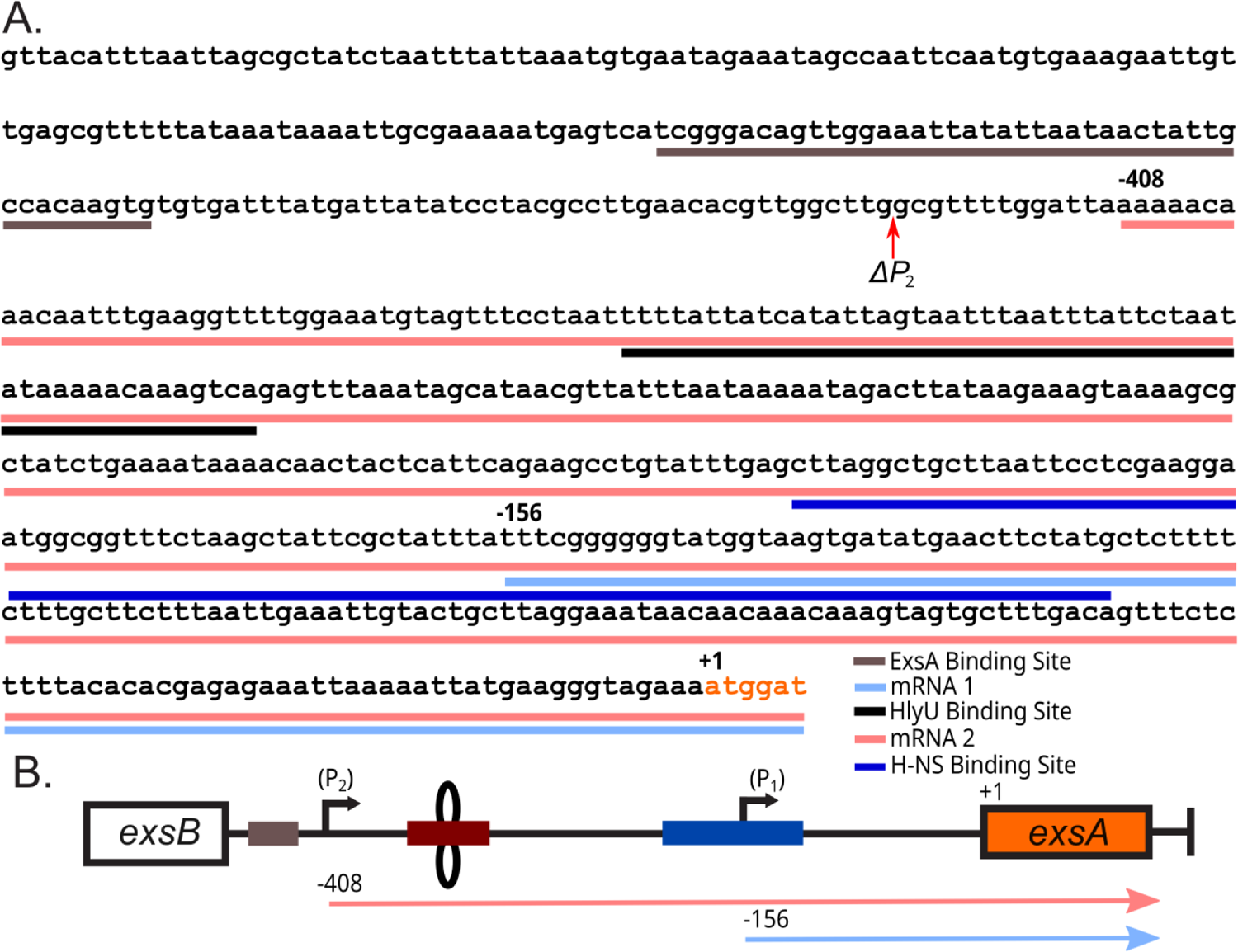
Two promoters exist at the *exsBA* intergenic region that transcribe *exsA.* (A) DNA sequence of the *exsBA* intergenic region with corresponding mRNAs labelled as discovered by 5’RACE analysis. The light blue line indicates the shorter mRNA1 species, which begins 156bp upstream of the *exsA* open reading frame, while the light red line refers to the longer mRNA2 species beginning 408bp upstream of the *exsA* open reading fram*e (exsA* ORF indicated by orange text). The ExsA protein binding site is indicated by the brown line^1, 2^, and the HlyU protected region previously reported (8) is indicated by the black line. The red arrow identifies the beginning of the **Δ**P_2_ DNA fragment used to verify the existence of the second promoter (Fig 6). (B) Schematic diagram of mRNA species identified via 5’RACE analysis in panel A. Colour scheme follows from panel A.

**Table S1.**
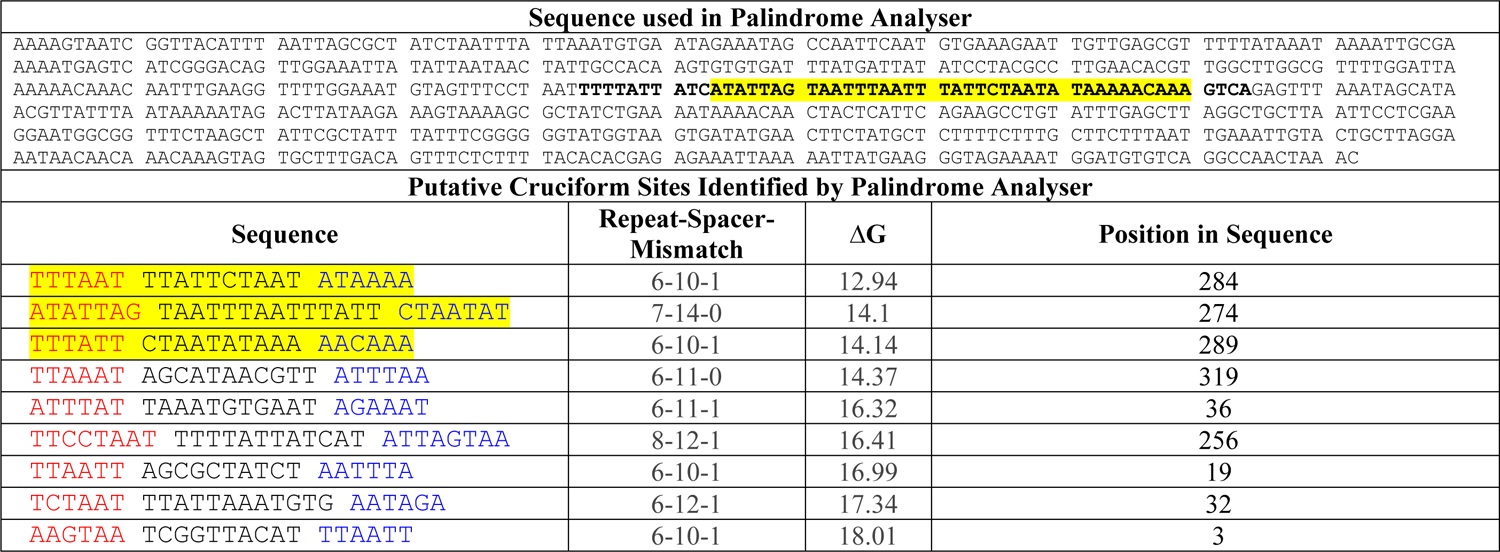
Intergenic sequence from *exsBA* in *Vibrio parahaemolyticus* was used as input into Palindrome Analyser -- an online bioinformatics tool which identifies putative cruciform forming sequences in provided nucleotide sequences and calculates the amount of energy required for cruciform formation. For each sequence, the 10 (or fewer) possible cruciforms requiring the smallest change in free energy to form are detailed. The HlyU protected region is bolded. Cruciform structures which overlap the HlyU protected region previously identified are highlighted.

**Table S2.**
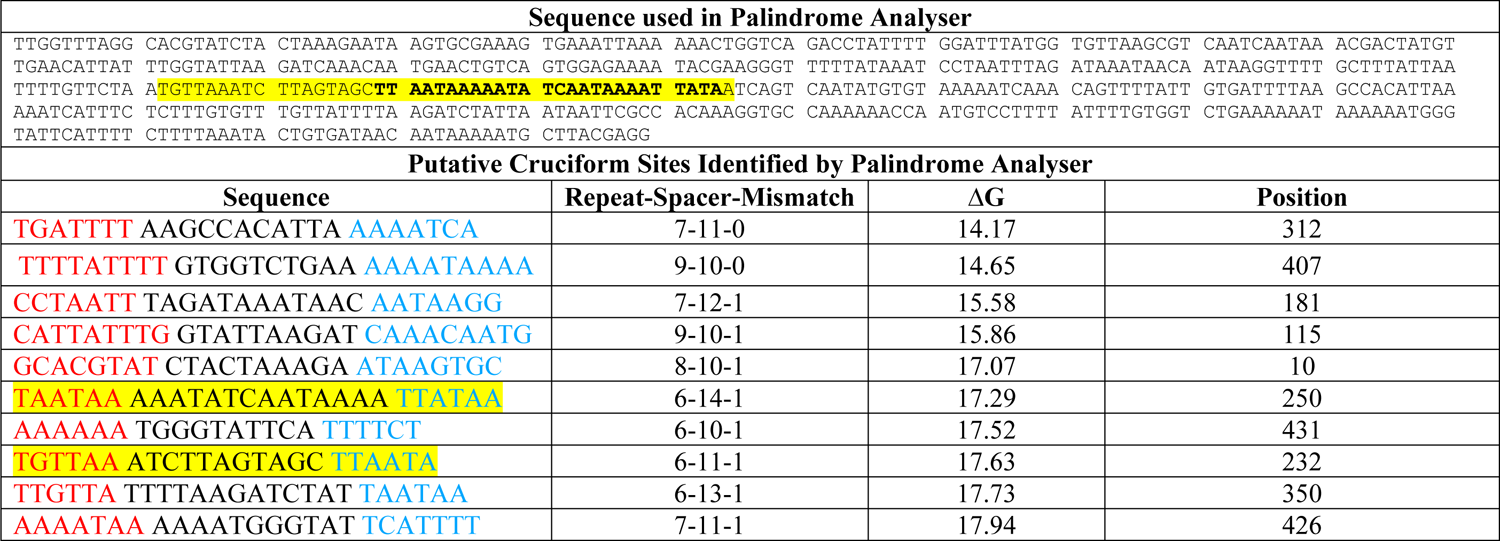
Intergenic sequence from *plp-vah* of *V. anguillarum* was used as input into Palindrome Analyser - an online bioinformatics tool which identifies putative cruciform forming sequences in provided nucleotide sequences and calculates the amount of energy required for cruciform formation. For each sequence, the 10 (or fewer) possible cruciforms requiring the smallest change in free energy to form are detailed. The DNA sequence in bold font represents the known HlyU binding site at this genetic locus. The HlyU protected region previously identified is bolded^3^. Cruciform structures which overlap the HlyU protected region previously identified are highlighted.

**Table S3.**
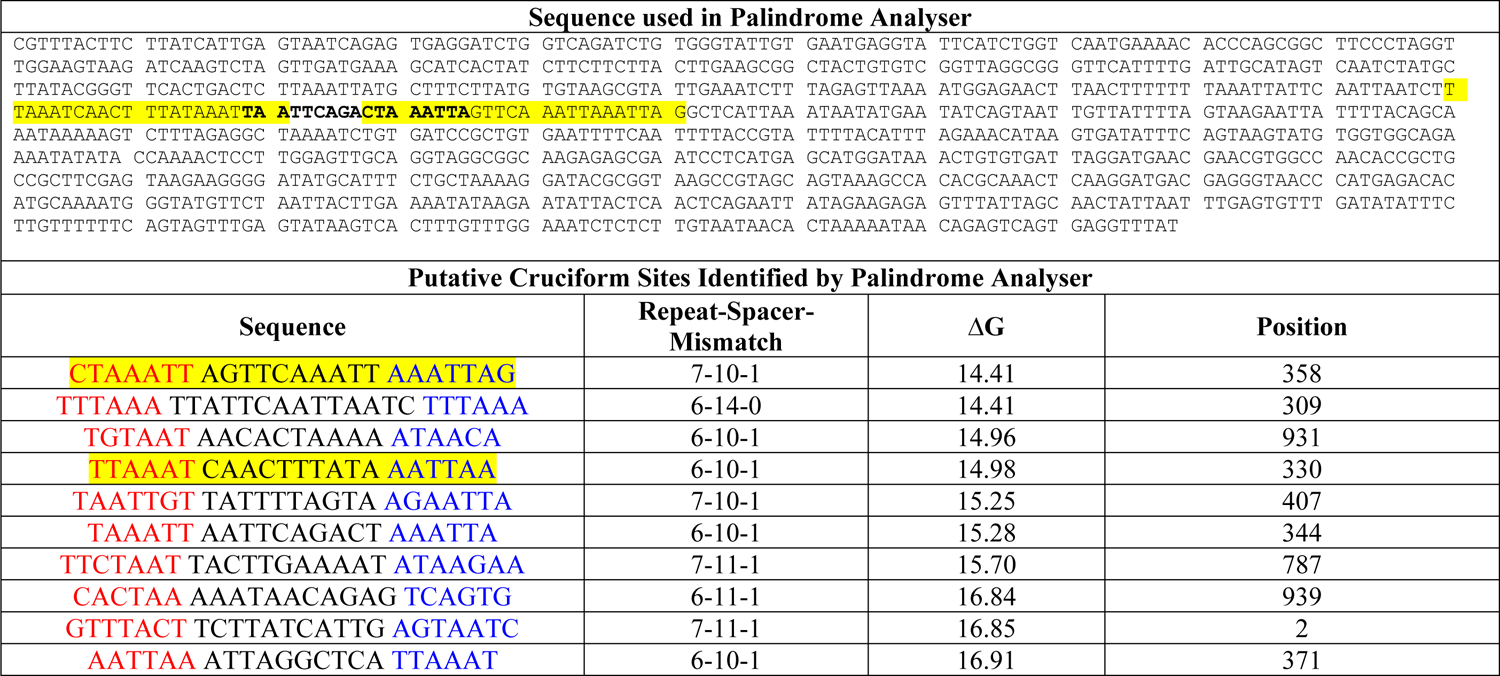
Intergenic sequence from *tlh-hlyA* of *V. cholerae* was used as input into Palindrome Analyser - an online bioinformatics tool which identifies putative cruciform forming sequences in provided nucleotide sequences and calculates the amount of energy required for cruciform formation. For each sequence, the 10 (or fewer) possible cruciforms requiring the smallest change in free energy to form are detailed. The lowest free energy cruciform is indicated as yellow highlighted text. The DNA sequence in bold font represents the known HlyU binding site at this genetic locus. The HlyU protected region previously identified is bolded^4^. Cruciform structures which overlap the HlyU protected region previously identified are highlighted.

**Table S4.**
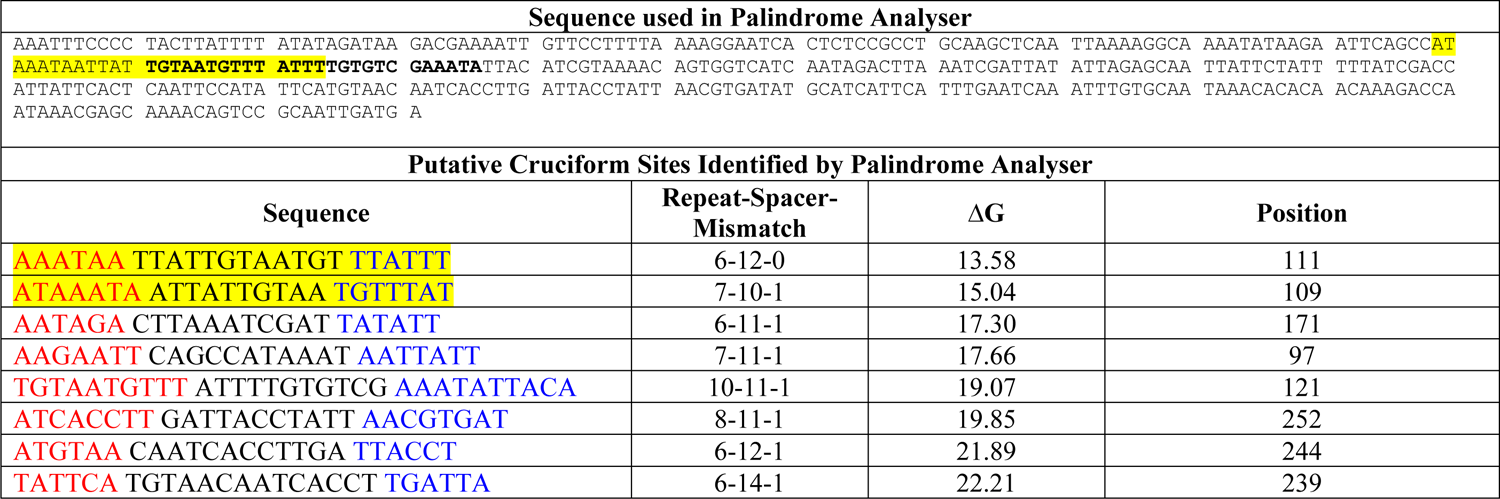
Intergenic sequence from the *rtxA1* operon of *V. vulnificus* was used as input into Palindrome Analyser - an online bioinformatics tool which identifies putative cruciform forming sequences in provided nucleotide sequences and calculates the amount of energy required for cruciform formation. For each sequence, the 10 (or fewer) possible cruciforms requiring the smallest change in free energy to form are detailed. The HlyU binding site is indicated in bold font. Cruciform structures which overlap the HlyU protected region previously identified are highlighted.

**Table S5.** Observed cell cytotoxicity of bacterial strains with or without complementation of *exsA*. +++: cytotoxicity equal to wildtype, ++: cytotoxicity less than wildtype, +: significantly less than wildtype, -: no cytotoxicity (relative to uninfected control). N=2

| Strain | Cytotoxicity Score |
| --- | --- |
| Wildtype/VSV105 (Positive) | +++ |
| $\Delta hlyU$/VSV105 (Negative) | - |
| WT/VSV105-*exsA* | +++ |
| $\Delta hlyU$/VSV105-*exsA* | +++ |
| IR1/VSV105-*exsA* | +++ |
| IR2/VSV105-*exsA* | +++ |
| PAL2/VSV105-*exsA* | + |
| IV1/VSV105-*exsA* | ++ |
| IV2/VSV105-*exsA* | +++ |
| IV3/VSV105-*exsA* | +++ |

**Table S6.** Table of bacterial strains used in this study.

| Strain Name | Organism | Genotype | Source |
| --- | --- | --- | --- |
| RIMD2210633 | *V. parahaemolyticus* | Wildtype | 1 |
| $\Delta hns$ | *V. parahaemolyticus* | *hns* null | This study |
| $\Delta vscN1$ | *V. parahaemolyticus* | *vscN1* null | 2 |
| $\Delta hlyU$ | *V. parahaemolyticus* | *hlyU* null | 3 |
| IR1 | *V. parahaemolyticus* | IR1 mutation in *exsA* upstream region | This study |
| IR2 | *V. parahaemolyticus* | IR2 mutation in *exsA* upstream region | This study |
| PAL2 | *V. parahaemolyticus* | PAL2 mutation in *exsA* upstream region | This study |
| IV1 | *V. parahaemolyticus* | IV1 mutation in *exsA* upstream region | This study |
| IV2 | *V. parahaemolyticus* | IV2 mutation in *exsA* upstream region | This study |
| IV3 | *V. parahaemolyticus* | IV3 mutation in *exsA* upstream region | This study |
| *tdh::exsBA-luxCDABE* | *V. parahaemolyticus* | *exsBA* intergenic region fused to promoter-less *luxCDABE* cassette and integrated into the *tdh* locus on chromosome | 3 |
| Tn5::*hns tdh::exsBA-luxCDABE* | *V. parahaemolyticus* | Tn5 insertion in *hns*. Strain harbors the *exsBA* intergenic region fused to promoter-less *luxCDABE* cassette integrated into the *tdh* locus on chromosome | 3 |
| DH5α*pir* | *E. coli* | | Stratagene |
| BL21(λDE3) | *E. coli* | | Novagen |

**Table S7.** Table of plasmids used in this study.

| Plasmid Name | Description | Source |
| --- | --- | --- |
| pVSV105 | Vibrio Shuttle Vector 105 contains *cat*, *oriT*, a *Vibrio* host *ori*, and *oriR6Kγ*. *lac* promoter. | 4 |
| pVSV105-*exsA* | VSV105 containing the *exsA* coding gene ahead of the *lac* promoter | This study |
| pEVS104 | Helper plasmid for tri-parental matings with *oriT* plasmids. | 5 |
| pVSVlux | pVSV105 containing *luxCDABE* cassette from pJW15 cloned blunt into the *Sma*I site opposite of the *lac* promoter. | 3 |
| pVSVlux-*exsBA* | pVSVlux containing the WT *exsBA* intergenic region (656 bp upstream of *exsA* start site) | 3 |
| pVSVlux-*exsBA*ΔIR | pVSVlux containing the *exsBA* intergenic region deleted for inverted repeat 1 | This study |
| pVSVlux-*exsBA*ΔIR2 | pVSVlux containing the *exsBA* intergenic region deleted for inverted repeat 2 | This study |
| pVSVlux-*exsBA*ΔIR1ΔIR2 | pVSVlux containing the *exsBA* intergenic region deleted for inverted repeats 1 and 2 | This study |
| pVSVlux-*exsBA*ΔPAL | pVSVlux containing the *exsBA* intergenic region deleted for central palindrome A/T rich sequence | This study |
| pVSVlux-IR1 | pVSVlux containing the IR1 mutation containing *exsBA* intergenic region | This study |
| pVSVlux-IR2 | pVSVlux containing the IR2 mutation containing *exsBA* intergenic region | This study |
| pVSVlux-PAL2 | pVSVlux containing the PAL2 mutation containing *exsBA* intergenic region | This study |
| pVSVlux-IV1 | pVSVlux containing the IV1 mutation containing *exsBA* intergenic region | This study |
| pVSVlux-IV2 | pVSVlux containing the IV2 mutation containing *exsBA* intergenic region | This study |
| pVSVlux-IV3 | pVSVlux containing the IV3 mutation containing *exsBA* intergenic region | This study |
| pFLAG-CTC-*hlyU* | pFLAG-CTC vector containing the open reading frame coding HlyU under the control of the *tac* promoter | 3 |
| pRK415-*exsA* | pRK415 vector containing the open reading frame coding ExsA under the control of the *tac* promoter | This study |
| pET21-*hlyU* | *hlyU* coding gene from *V. parahaemolyticus* cloned in frame with a C-terminal 6x HIS tag. IPTG inducible expression. | 3 |
| pRE112 | Allelic exchange vector. Contains *cat* and *sacB* for allelic exchange procedure and *oriR6Kγ* origin (suicide vector). | 6 |
| pRE112-IR1 | Allelic exchange plasmid pRE112 containing the IR1 mutation in the *exsBA* intergenic region. | This study |
| pRE112-IR2 | Allelic exchange plasmid pRE112 containing the IR2 mutation in the *exsBA* intergenic region. | This study |
| pRE112-PAL2 | Allelic exchange plasmid pRE112 containing the PAL2 mutation in the *exsBA* intergenic region. | This study |
| pRE112-IV1 | Allelic exchange plasmid pRE112 containing the IV1 mutation in the *exsBA* intergenic region. | This study |
| pRE112-IV2 | Allelic exchange plasmid pRE112 containing the IV2 mutation in the *exsBA* intergenic region. | This study |
| pRE112-IV3 | Allelic exchange plasmid pRE112 containing the IV3 mutation in the *exsBA* intergenic region. | This study |
| pUC(AT) | Positive control cruciform construct for T7 endonuclease assays | NEB |
| pBluescript | General cloning vector, plasmid for cloning gBlock DNA and for use as negative control in T7 endonuclease assays | Stratagene |
| p*exsBA*/pBS | PCR amplified *V. parahaemolyticus exsBA* intergenic region cloned into pBluescript for use in T7 endonuclease assays | This study |
| p*hlyA*/pBS | Synthetic gene block for *V. cholerae hlyA* region cloned into pBluescript for use in T7 endonuclease assays | This study |
| p*rtxA1*/pBS | Synthetic gene block for *V. vulnificus rtxA1* region cloned into pBluescript for use in T7 endonuclease assays | This study |
| p*rtxHB*/pBS | Synthetic gene block for *V. anguillarum rtxHB* region cloned into pBluescript for use in T7 endonuclease assays | This study |
| pVSVlux-$\Delta P_2$ | $\Delta P_2$ deletion of the *exsBA* intergenic region cloned into pVSVlux for studying $P_1$ expression in isolation | This study |

**Table S8.**
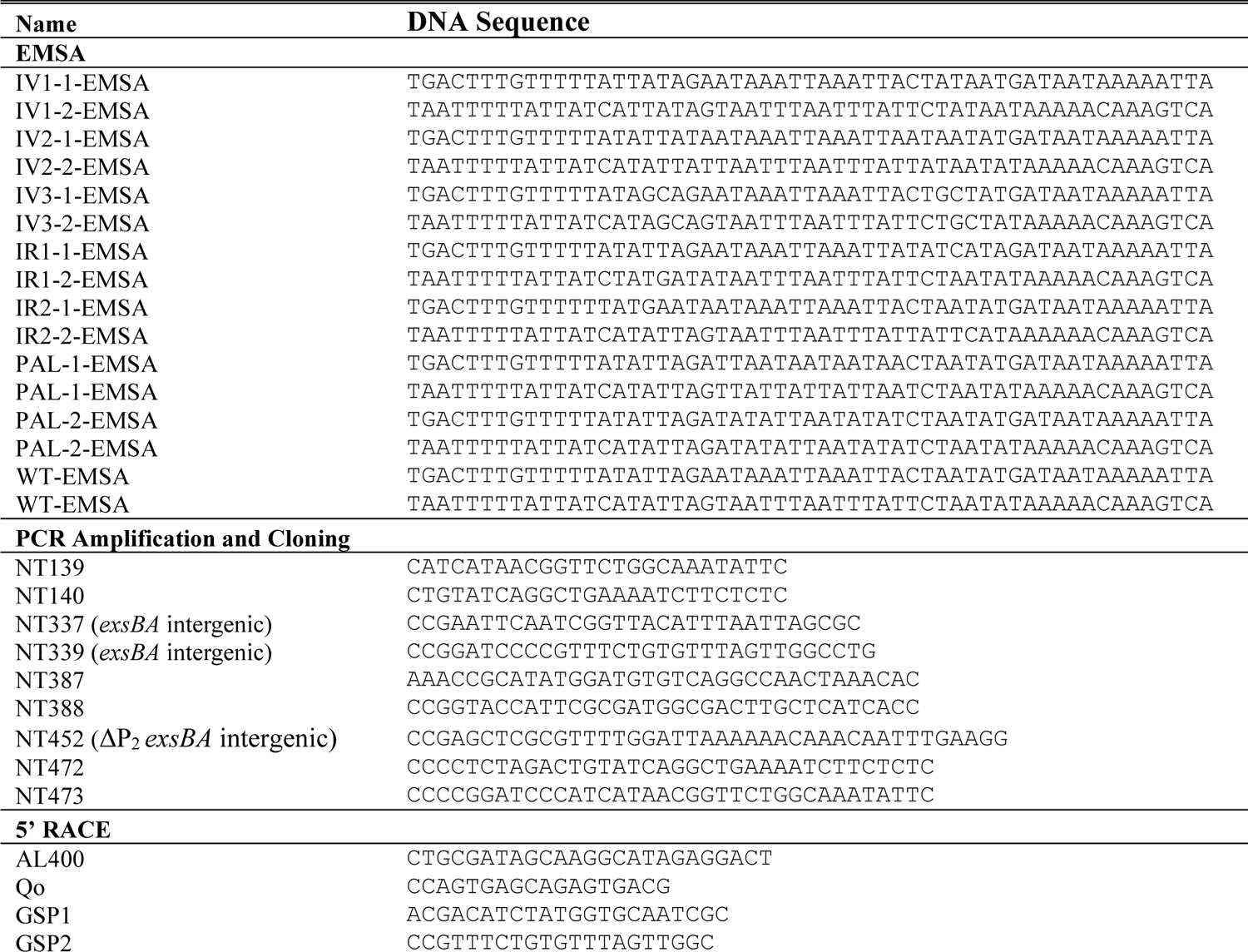
Table of oligonucleotides used in this study.

**Table S9.**
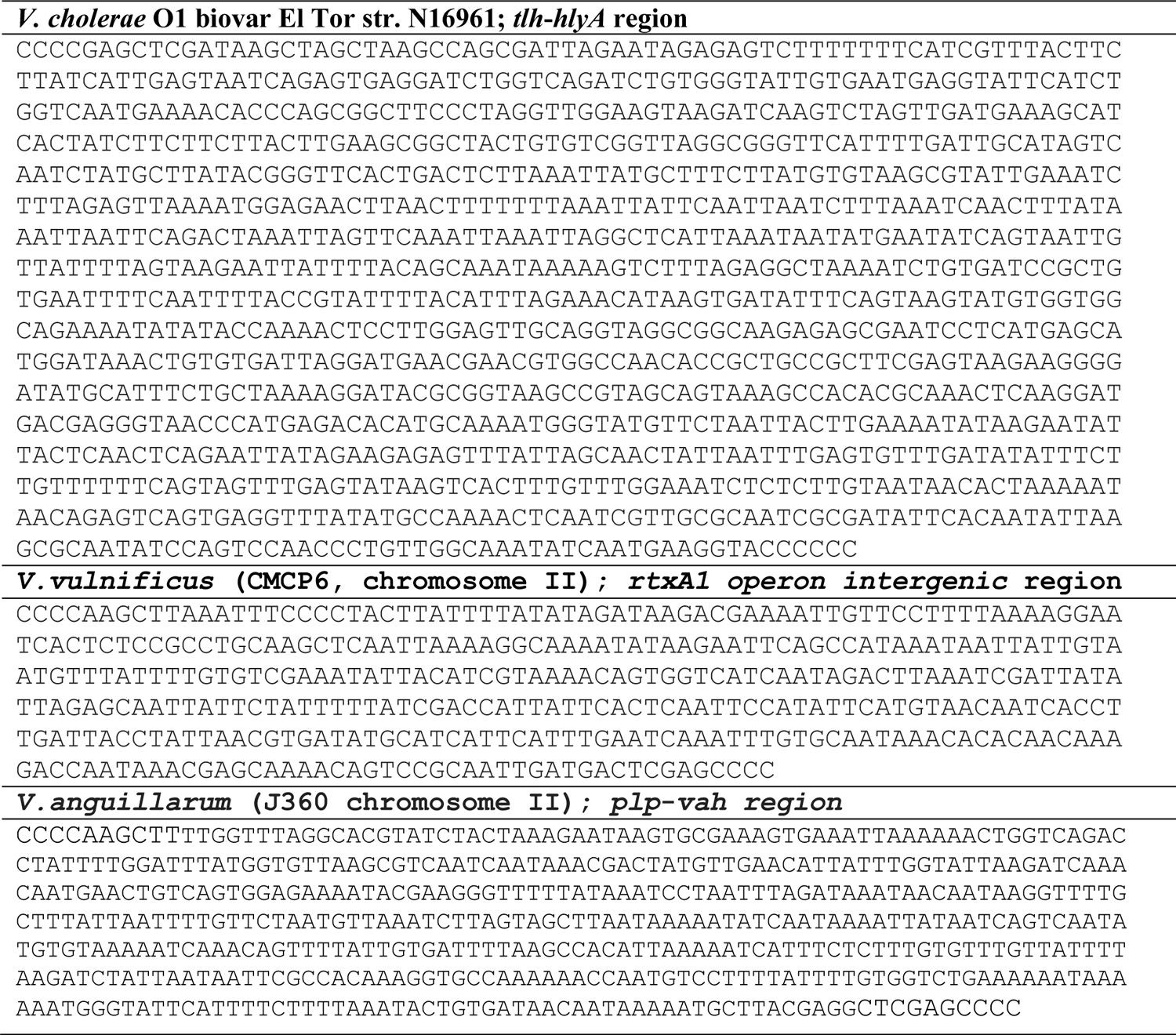
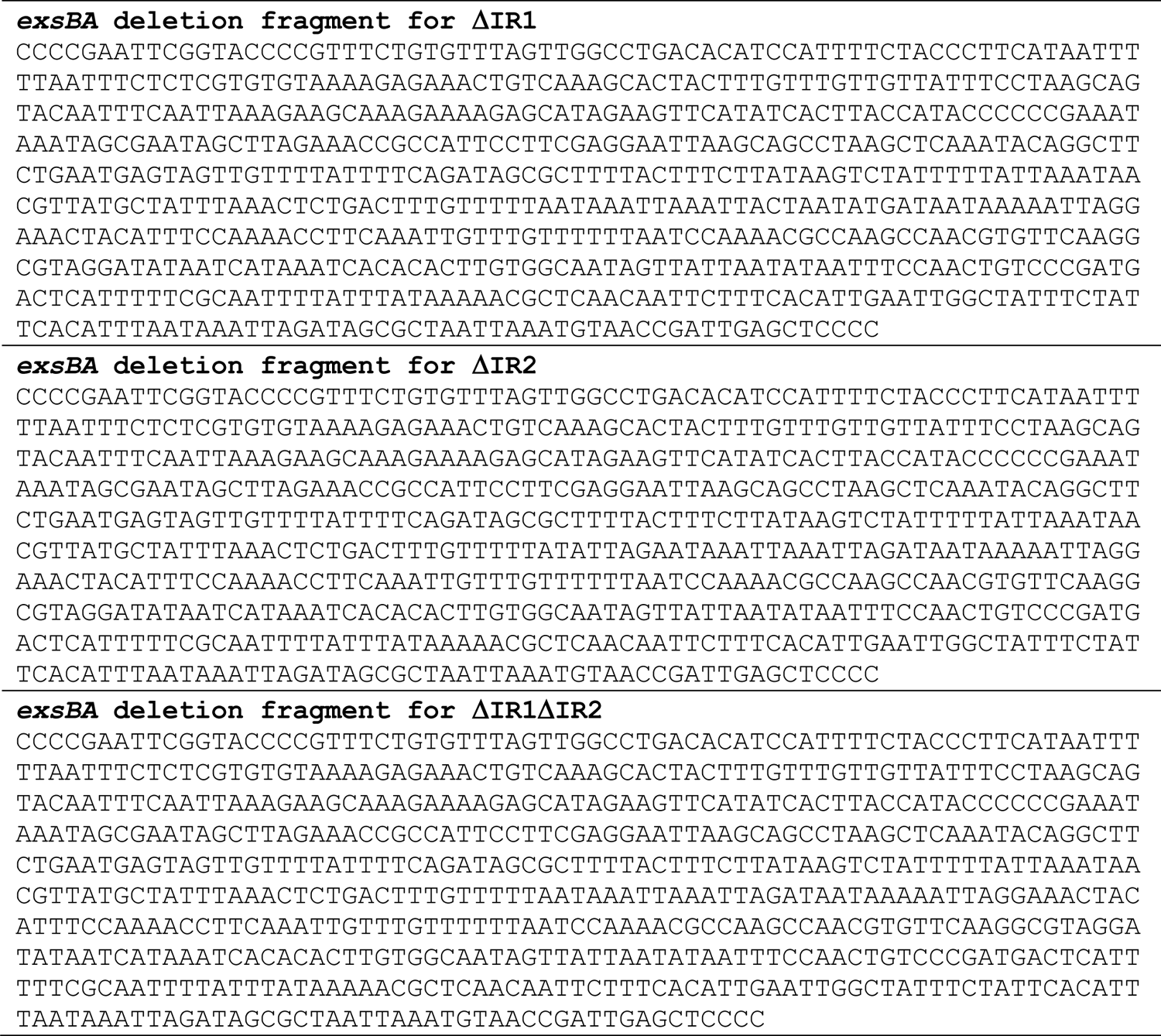
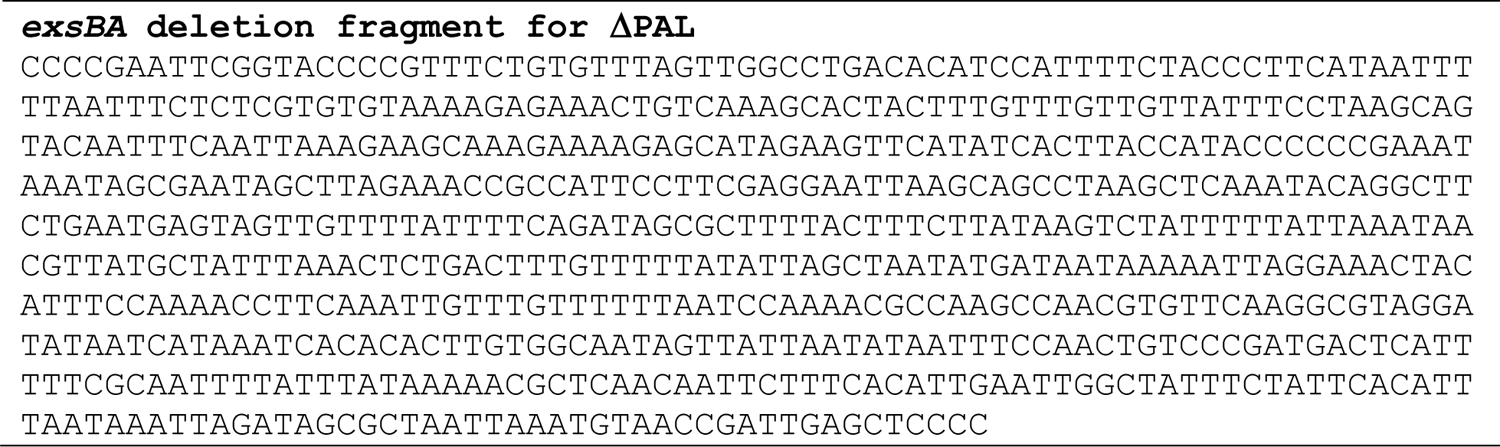
Synthetic gBlock DNA sequences for respective *Vibrio* spp in T7 endonuclease assays and *exsBA* intergenic deletion analyses.

